# Statistically consistent divide-and-conquer pipelines for phylogeny estimation using NJMerge

**DOI:** 10.1101/469130

**Authors:** Erin K. Molloy, Tandy Warnow

## Abstract

**Background:** Divide-and-conquer methods, which divide the species set into overlapping subsets, construct a tree on each subset, and then combine the subset trees using a supertree method, provide a key algorithmic framework for boosting the scalability of phylogeny estimation methods to large datasets. Yet the use of supertree methods, which typically attempt to solve NP-hard optimization problems, limits the scalability of such approaches.

**Results:** In this paper, we introduce a divide-and-conquer approach that does not require supertree estimation: we divide the species set into pairwise disjoint subsets, construct a tree on each subset using a base method, and then combine the subset trees using a distance matrix. For this merger step, we present a new method, called NJMerge, which is a polynomial-time extension of Neighbor Joining (NJ); thus, NJMerge can be viewed either as a method for improving traditional NJ or as a method for scaling the base method to larger datasets. We prove that NJMerge can be used to create divide-and-conquer pipelines that are statistically consistent under some models of evolution. We also report the results of an extensive simulation study evaluating NJMerge on multi-locus datasets with up to 1000 species. We found that NJMerge sometimes improved the accuracy of traditional NJ and substantially reduced the running time of three popular species tree methods (ASTRAL-III, SVDquartets, and “concatenation” using RAxML) without sacrificing accuracy. Finally, although NJMerge can fail to return a tree, in our experiments, NJMerge failed on only 11 out of 2560 test cases.

**Conclusions:** Theoretical and empirical results suggest that NJMerge is a valuable technique for large-scale phylogeny estimation, especially when computational resources are limited. NJMerge is freely available on Github (http://github.com/ekmolloy/njmerge).

## Introduction

Estimating evolutionary trees, called phylogenies, from molecular sequence data is a fundamental problem in computational biology, and building the Tree of Life is a scientific grand challenge. It is also a computational grand challenge, as many of the most accurate phylogeny estimation methods are heuristics for NP-hard optimization problems. Species tree estimation can be further complicated by biological processes (e.g., incomplete lineage sorting, gene duplication and loss, and horizontal gene transfer) that create heterogeneous evolutionary histories across genomes or “gene tree discordance” [24].

Incomplete lineage sorting, which is modeled by the Multi-Species Coalescent (MSC) model [36, 39], has been shown to present challenges for phylogenomic analyses [11]. In addition, while the standard approach for multi-locus species tree estimation uses maximum likelihood methods (e.g., RAxML) on the concatenated multiple sequence alignment, recent studies have established that even exact algorithms for maximum likelihood are not statistically consistent methods for multi-locus species tree estimation under the MSC model (see [42] for a proof for unpartitioned maximum likelihood and [41] for fully partitioned maximum likelihood).

Because concatenation analyses using maximum likelihood are provably not statistically consistent in the presence of incomplete lineage sorting, new methods have been developed that are provably statistically consistent under the MSC model. Bayesian methods that coestimate gene trees and species trees (e.g., [15, 35]) are statistically consistent and expected to be the highly accurate; however, such methods are also prohibitively expensive on large datasets. More efficient approaches have been developed that are statistically consistent under the MSC model, including “gene tree summary methods”, which take a collection of gene trees as input and then compute a species tree from the gene trees using only the gene tree topologies. For example, NJst [23] runs Neighbor Joining (NJ) [43] on the “average gene tree internode distance” (AGID) matrix, and ASTRAL [27] finds a quartet-median tree (i.e. a species tree that maximizes the total quartet tree similarity to the input gene trees) within a constrained search space. However, gene tree summary methods can have reduced accuracy when gene tree estimation error is high, which is a problem for many phylogenomic datasets (see discussion in [31]).

Because of the impact of gene tree estimation error, alternative approaches that bypass gene tree estimation, called “site-based” methods, have been proposed. Perhaps the best known of site-based method is SVDquartets [9], which estimates quartet trees from the concatenated sequence alignments (using statistical properties of the MSC model and the sequence evolution model) and then combines the quartet trees into a tree on the full set of species using quartet amalgamation methods that are heuristics for the Maximum Quartet Consistency problem [18]. Other examples of site-based methods include computing JukesCantor [20] or log-det [46] distances from concatenated alignment and then running NJ on the resulting distance matrix. Such approaches can be statistically consistent under the MSC model when the sequence evolution models across genes satisfy some additional assumptions (e.g., a relaxed molecular clock) [10, 3].

Many of these methods (e.g., ASTRAL, SVDquartets, and concatenation using RAxML) are heuristics for NP-hard optimization problems. Such methods can have difficulties scaling to datasets with large numbers of species, and divide-and-conquer approaches have been developed to scale methods to larger datasets (e.g., the family of disk covering methods [55, 16, 21, 32, 6, 53). Such methods operate by dividing the species set into overlapping subsets, constructing trees on the subsets, and then merging the subset trees into a tree on the entire species set. The last step of this process, called “supertree estimation”, can provide good accuracy (i.e., retain much of the accuracy in the subset trees) if good supertree methods are used. Notably, the supertree compatibility problem is NP-complete [7], and the preferred supertree methods attempt to solve NP-hard optimization problems (e.g., the Robinson-Foulds supertree problem [5], the Maximum Quartet Consistency problem [19], the Matrix Representation with Parsimony problem [38], and the Matrix Representation with Likelihood problem [34]). In summary, none of the current supertree methods provide both accuracy and scalability to datasets with large numbers of species (see [54] for further discussion).

In this paper, we introduce a new divide-and-conquer approach to scaling phylogeny estimation methods to large datasets: we divide the species (or leaf) set into pairwise disjoint subsets, construct a tree on each of the subsets, and then assemble the subset trees into a tree on the entire species set. Supertree methods cannot be used to combine trees on pairwise disjoint leaf sets, and we present a new polynomial-time method, called NJMerge, for this task. We prove that NJMerge can be used in statistically consistent divide-and-conquer pipelines for both gene tree and species tree estimation and evaluate the effectiveness of using NJMerge in the context of multi-locus species tree estimation. We found, using an extensive simulation study, that NJMerge sometimes improved the accuracy of traditional NJ and that NJMerge provided substantial improvements in the running time for three methods (ASTRAL-III [57], SVDquartets [9], and concatenation using RAxML [44]) without sacrificing accuracy. Furthermore, NJMerge enabled SVDquartets and RAxML to run on large datasets (e.g., 1000 taxa and 1000 genes), on which SVDquartets and RAxML would otherwise fail to run when limited to 64 GB of memory. While NJMerge is not guaranteed to return a tree; the failure rate in our experiments was low (less than 1% of tests). In addition, NJMerge failed on fewer datasets than either ASTRAL-III, SVDquartets, or RAxML — when given the same computational resources: a single compute node with 64GB of physical memory, 16 cores, and a maximum wall-clock time of 48 hours. Together, these results suggest that NJMerge is a valuable technique for large-scale phylogeny estimation, especially when computational resources are limited.

## NJMerge

Neighbor Joining (NJ) [43], perhaps the most widely used polynomial-time method for phylogeny estimation, estimates a tree *T* from a dissimilarity matrix *D*; NJMerge is a polynomial-time extension of NJ to impose a set of constraints on the output tree *T* (Figure 1). More formally, NJMerge takes as input a dissimilarity matrix *D* on leaf set *S* = {*s*_1_, *s*_2_, …, *s*_*n*_} and a set 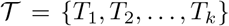 of unrooted binary trees on pairwise disjoint subsets of the leaf set *S* and returns a tree *T* that agrees with every tree in 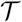 (Definition 1). Note that the output tree *T* is a compatibility supertree for 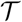 and that because the trees in 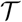 are on pairwise disjoint subsets of the leaf set *S*, a compatibility supertree always exists. NJMerge does not require that the input constraint trees 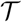 to form clades in *T*. For example, the caterpillar tree on **{***A, B, C, D, E, F, G, H***}** obtained by making a path with the leaves hanging off it in alphabetical order is a compatibility supertree for 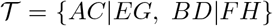, and yet the trees in 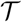 do not form clades within the caterpillar tree (Figure 2). Of course, other compatibility supertrees exist for 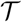, and, in some of them, the input constraint trees will form clades. The objective is to find a tree that is close to the true (but unknown) tree from the set of all compatibility supertrees for 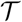, and NJMerge tries to achieve this objective by using the dissimilarity matrix *D*.

**Figure 1:**
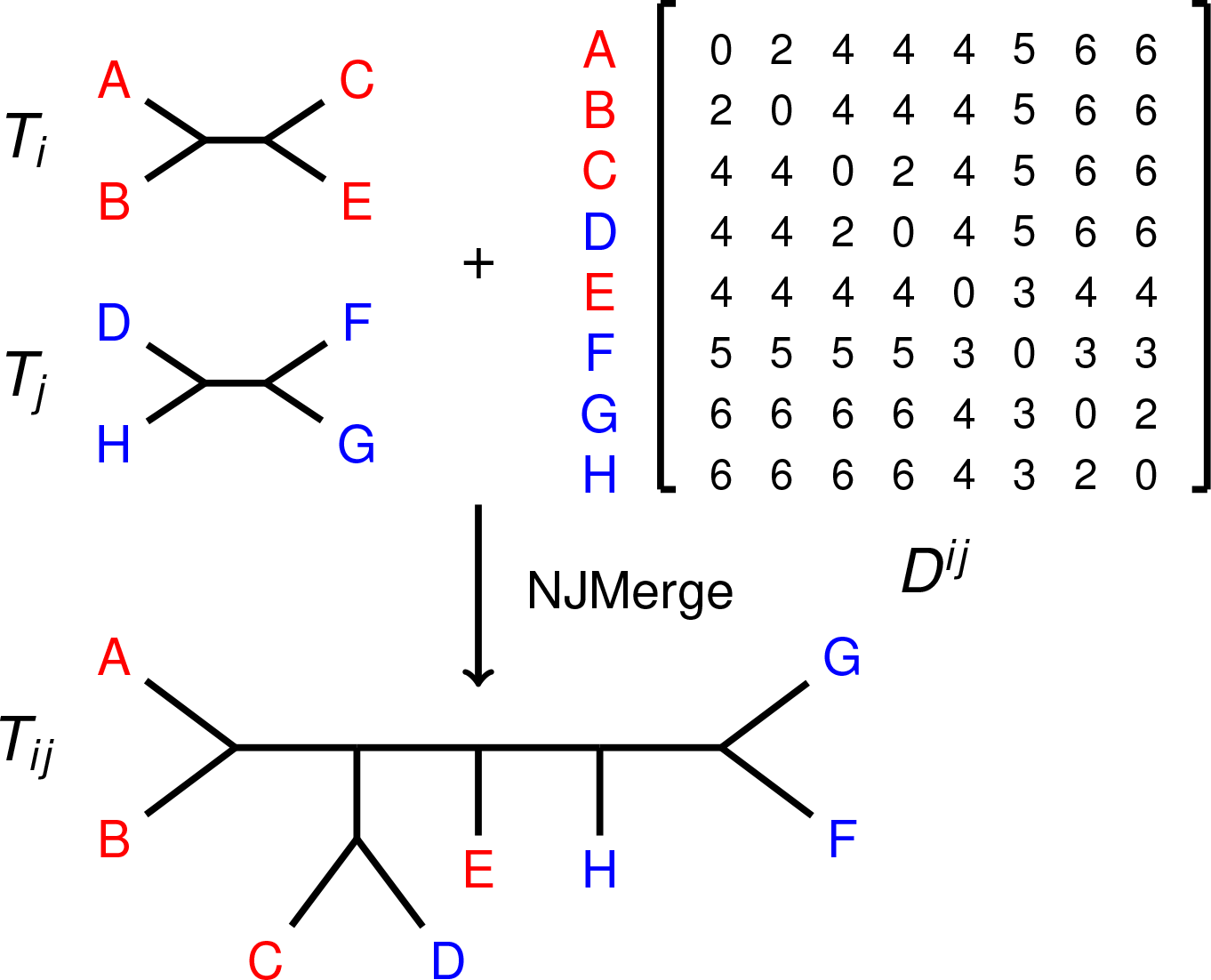
NJMerge Input/Output Example. In this example, NJMerge is given two constraint trees (*T*_*i*_ and *T*_*j*_) and a distance matrix *D*^*ij*^ that is additive for the tree (((*A, B*), (*C, D*)), *E*, (*F,* (*G, H*))). NJMerge returns a compatibility supertree, called *T*_*ij*_, for the two constraint trees (*T*_*i*_ and *T*_*j*_). Note that Neighbor Joining (NJ) applied to the distance matrix *Dij* would return ((((*A, B*), (*C, D*)), *E*, (*F,* (*G, H*))) [4]; however, NJMerge rejects the siblinghood proposal (*G, H*), because it violates constraint tree *T*_*j*_. Instead, NJMerge makes *G* and *F* siblings.

**Figure 2:**
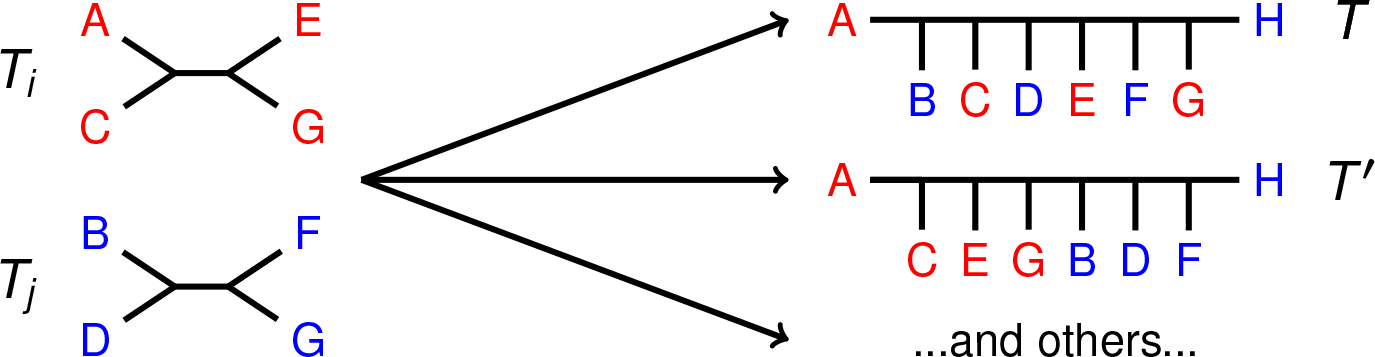
Compatibility Supertree Example. In this example, two compatibility supertrees for 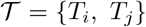 are shown. Note that the trees in 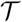 form clades in *T*′ but do not form clades in *T*. Other compatibility supertrees for 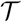 exist.

### Definition 1.

*Let T be a tree on leaf set S, and let T′ be a tree on leaf set R ⊆ S. We say that T′* **agrees with** *T if restricting T to leaf set R induces a binary tree that (after suppressing the internal nodes of degree 2) is isomorphic to T′*.

Here we briefly describe the NJ algorithm by Saitou and Nei [43]. NJ has an iterative design that builds the tree from the bottom up, producing a rooted tree that is then unrooted. Initially, all *n* leaves are in separate components. When a pair of leaves is selected to be siblings, the pair of leaves is effectively replaced by a rooted tree on two leaves, and the number of components is reduced by one. This process repeats until there is only one component: a tree on the full leaf set. At each iteration, NJ updates *D* based on the new sibling pair, derives a new matrix *Q* from *D*, and uses *Q* to determine which pair of the remaining nodes to join. Specifically, NJ accepts siblinghood proposal (*i, j*) such that *Q*[*i, j*] is minimized. The same formulas used by NJ [43] to update *D* and compute *Q* are also used by NJMerge; however, NJMerge can make different siblinghood decisions than NJ — based on the input constraint trees.

After each siblinghood decision, NJMerge updates the constraint trees. Specifically, when two leaves are made siblings, they are replaced by a new leaf, and the constraint trees are relabeled. For example, if *x* is a leaf in *T*_*i*_ and *y* is a leaf in *T*_*j*_, then the siblinghood proposal *z* = (*x, y*) requires that *x* and *y* are replaced with *z* in *T*_*i*_ and *T*_*j*_, respectively. Because siblinghood decisions change the set of leaves in the constraint trees, they can result in the constraint trees no longer being disjoint (Figure 3). Thus, siblinghood decisions have the potential to make the set of constraint trees incompatible. Determining whether or not a set of unrooted phylogenetic trees is compatible is an NP-complete problem [45, 52], so NJMerge uses a polynomial-time heuristic. In each iteration, NJMerge sorts the entries of the *Q* from least to greatest and accepts the first siblinghood proposal (*x, y*) that satisfies the following properties:

1. If *x* and *y* are both in some constraint tree *T*_*i*_, then they are siblings in *T*_*i*_.
2. If *x* or *y* are in more than one constraint trees, then replacing *x* and *y* with a new leaf *z* = (*x, y*) in all constraint trees does not make any *pair* of constraint trees incompatible, i.e., a compatibility supertree exists for every *pair* of updated constraint trees.

**Figure 3:**
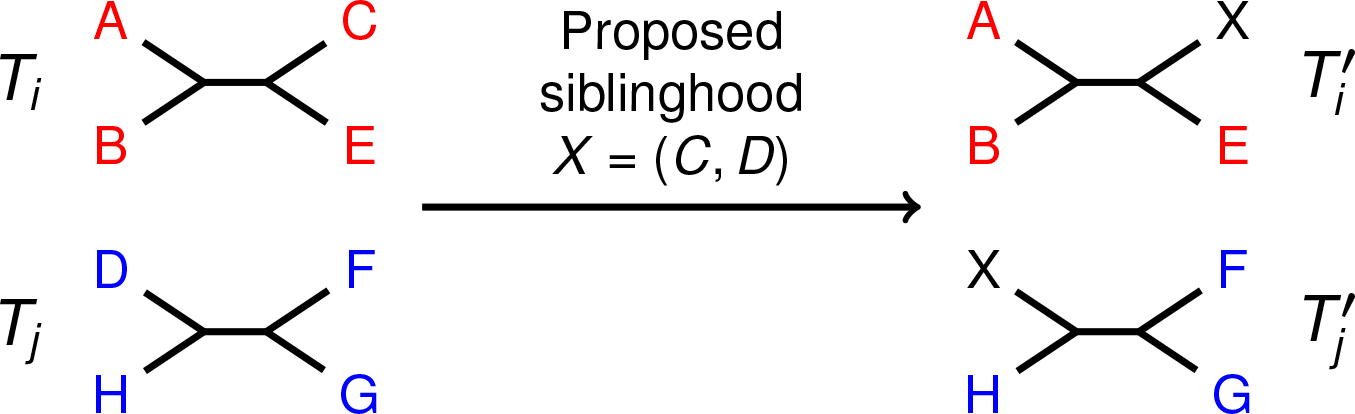
NJMerge Siblinghood Proposal Example. In this example, NJMerge evaluates the siblinghood proposal (*C, D*). Because *C* ∈ *T*_*i*_ and *D* ∈ *T*_*j*_, NJMerge first updates the constraint trees *T*_*i*_ and *T*_*j*_ based on the proposed siblinghood to get 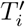 and 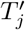. Specifically, both *C* ∈ *T*_*i*_ and *D* ∈ *T*_*j*_ are replaced by *X*, representing the siblinghood (*C, D*). The compatibility of the updated constraint trees can be tested by rooting the trees at leaf *X* and using the algorithm proposed in [1]. Because the updated constraint trees (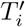 and 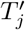) are indeed compatible, NJMerge will accept siblinghood proposal. Importantly, when NJMerge evaluates the next siblinghood proposal, the two constraint trees will no longer be on disjoint leaf sets.

Because pairwise compatibility of unrooted trees does not guarantee that the entire set of constraint trees is compatible, it is possible for NJMerge to accept a siblinghood decision that will eventually cause the algorithm to fail when none of the remaining leaves can be joined without violating the pairwise compatibility of constraint trees. Although the “pairwise compatibility heuristic” can fail, it is easy to see that if NJMerge returns a tree, then it is a compatibility supertree for the input set 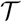 of constraint trees.

To determine if some pair of constraint trees becomes incompatible after making *x* and *y* siblings, it suffices to check only those pairs of constraint trees that contain at least one of *x* and *y*; all other pairs of trees are unchanged by accepting the siblinghood proposal and are pairwise compatible by induction. Because the leaves in the two trees labeled *x* or *y* have been relabeled by the new leaf *z* = (*x, y*), they can be treated as rooted trees by rooting them at *z*. Testing the compatibility of rooted trees is easily accomplished in polynomial time using [1]. In fact, instead of testing pairs of constraint trees, the entire set of trees in 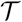 containing the new leaf *z* = (*x, y*) can be tested for compatibility in polynomial time using [1]. Furthermore, if at least one leaf exists in all constraint trees, then the compatibility of 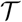 can be determined in polynomial time. Finally, note the input matrix was referred to as a dissimilarity matrix (and not a distance matrix), because estimated distances between species may not satisfy the triangle inequality [53]; however, this matrix is more commonly referred to as a distance matrix, and we use this term henceforth.

### Divide-and-Conquer Pipelines for Phylogeny Estimation

NJMerge can be used in divide-and-conquer pipelines for phylogeny estimation as shown in Figure 4 and described below. In order to run this pipeline, the user must select a method for decomposing the leaf set into pairwise disjoint subsets (step 2), a maximum subset size (step 2), a method for computing a distance matrix *M*_*D*_ (step 1), and a method *M*_*T*_ for computing subset trees (step 3); thus, the user can select *M*_*D*_ and *M*_*T*_ to be appropriate for gene tree estimation or species tree estimation. The pipeline then operates as follows.

1. Estimate distances between pairs of leaves using method *M*_*D*_.
2. Decompose the leaf set into pairwise disjoint subsets.

2a. Compute a starting tree by running NJ on the distance matrix computed in Step 1.
2b. Decompose the starting tree into pairwise disjoint subsets of leaves with a predefined maximum subset size (e.g., using the centroid tree decomposition described in PASTA [26]).
3. Build a tree on each subset using method *M*_*T*_, thus producing the set 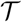 of constraint trees. Note that constraint trees can be estimated in serial or in parallel, depending on the computational resources available.
4. Run NJMerge on the input pair 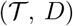.

**Figure 4:**
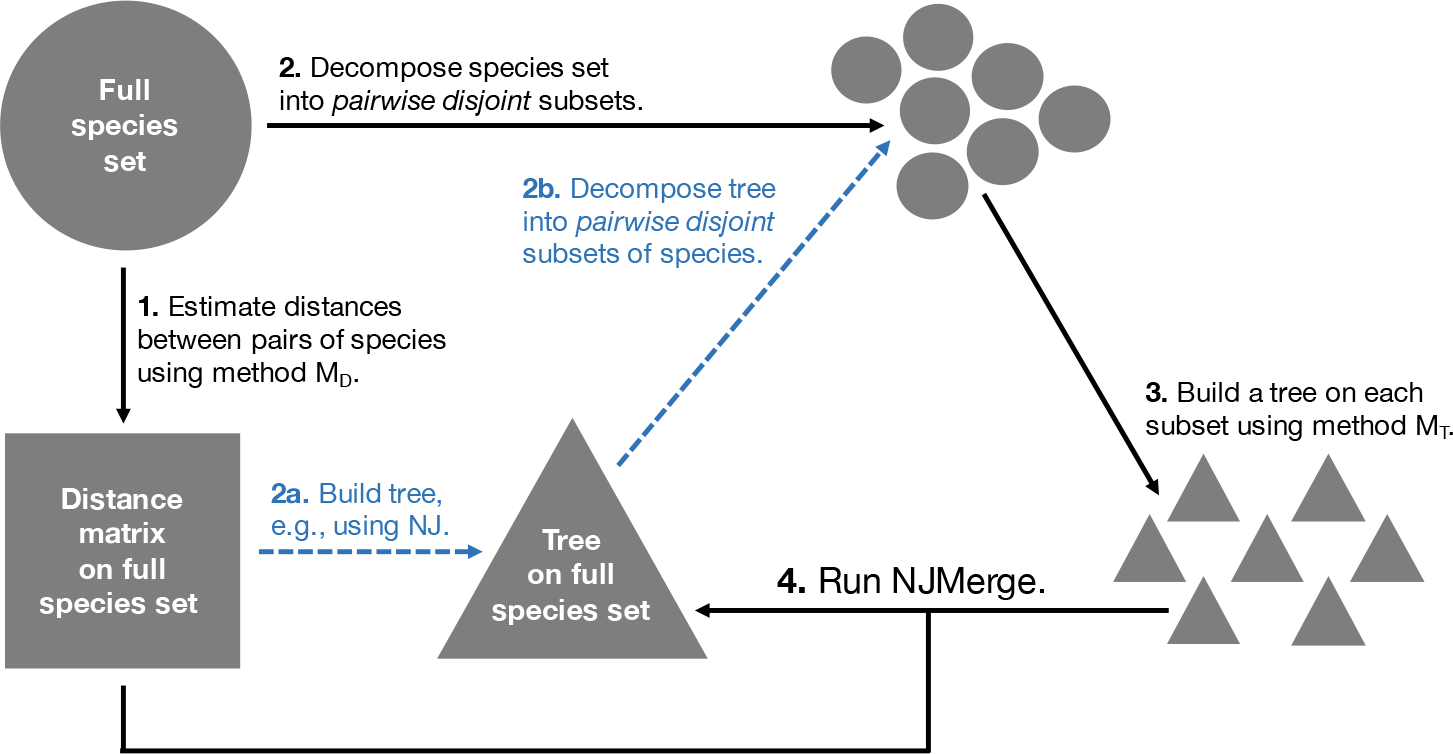
Divide-and-Conquer Pipeline using NJMerge. We present a divide-and-conquer pipeline that operates by 1) estimating distances between pairs of species using method *M*_*D*_, 2) decomposing the species set into pairwise disjoint subsets, 3) building a tree on each subset using method *M*_*T*_, and 4) merging trees together using the distance matrix using NJMerge. Step 2 can be performed by estimating a tree from the distance matrix (e.g., using NJ) and then decomposing this tree into pairwise disjoint subsets of species (shown in blue). Although not explored in this study, this pipeline can be run in an iterative fashion by using the tree produced in Step 4 to define the next subset decomposition. In this schematic, sets of species are represented by circles, distance matrices are represented by squares, and trees are represented by triangles.

Finally, although not explored in this study, this pipeline can be run in an iterative fashion by using the tree produced in step 4 to define the next subset decomposition.

### Statistical Consistency

Neighbor Joining (NJ) has been proven to be statistically consistent [14, 4, 8] under models of evolution for which pairwise distances can be estimated in a statistically consistent manner. This includes standard models of sequence evolution (e.g., the Generalized Time Reversible (GTR) model [50], which contains other models of sequence evolution, including Jukes-Cantor [20]). More recently, NJ has been used on multi-locus datasets to estimate species trees under the Multi-Species Coalescent (MSC) model; specifically, the method, NJst [23] estimates a species tree by running NJ on the average gene tree internode distance (AGID) matrix, calculated by averaging the topological distances between pairs of species in the input set of gene trees. Allman et al. [2] showed that the AGID matrix converges to an additive matrix for the species tree, and so NJst and some other methods (e.g., ASTRID [51]) that estimate species trees from the AGID matrix are statistically consistent under the MSC model.

We now prove that NJMerge can be used in statistically consistent divide-and-conquer pipelines for estimating gene trees and species trees. These results follow from Theorem 3 that shows NJMerge will return the tree *T*\* when given a nearly additive distance matrix (Definition 2) for *T*\* and a set 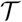 of constraint trees that agree with *T*\* (Definition 1).

#### Definition 2.

*Let T be a tree with positive weights on the edges and leaves labelled* 1, 2, …, *n*. *We say that an n* × *n matrix M is* **nearly additive** *for T if each entry M* [*i, j*] *differs from the distance between leaf i and leaf j in T by less than one half of the shortest branch length in T*.

#### Theorem 3.

*Let* 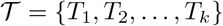 *be a set of trees, and let D be a distance matrix on S* = ⋃_*i*_ *S*_*i*_, *where S*_*i*_ *is the set of leaves in T*_*i*_. *Let T*\* *be a tree on leaf set S*. *If D is a nearly additive matrix for T*\* *and if T*_*i*_ *agrees with T*\* *for all i* ∈ {1,…, *k*}, *then NJMerge applied to input* 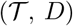 *returns T*\*.

*Proof.* NJ applied to a nearly additive distance matrix for *T*\* will return *T*\* [4]. Because all trees in 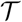 agree with *T*\*, the siblinghood proposals suggested by NJ will never violate the trees in 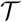 or the compatibility of 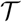. Thus, NJMerge applied to 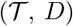 will return the same output as NJ applied to *D*, which is *T*\*.

We now define statistical consistency in the context of gene tree estimation (Definition and show that NJMerge can be used to create statistically consistent divide-and-conquer pipelines for gene tree estimation (Corollary 5).

#### Definition 4.

*Let* (*T,* Θ) *be a GTR model tree with topology T and numerical parameters* Θ *(e.g., substitution rate matrix, branch lengths, etc). A method M for constructing gene trees from DNA sequences is* **statistically consistent** *under the GTR model if, for all *ϵ* > 0, there exists a constant l* > 0 *such that, given sequences of length at least l, M returns T with probability at least* 1 − *ϵ*.

#### Corollary 5.

*NJMerge can be used in a gene tree estimation pipeline that is statistically consistent under the GTR model of sequence evolution*.

*Proof*. Let (*T*\*, Θ) be a GTR model tree, let *M*_*D*_ be a method for calculating distances between pairs of sequences, and let *M*_*T*_ be a method for constructing trees from DNA sequences. Suppose that

- the divide-and-conquer pipeline produces *k* pairwise disjoint subsets of sequences
- Neighbor Joining (NJ) applied to a matrix of pairwise distances calculated using *M*_*D*_ is a statistically consistent method for constructing gene trees under the GTR model (e.g., the log-det distance [46])
- *M*_*T*_ is statistically consistent under the GTR model (e.g., maximum likelihood [33, 12])

Now let *ϵ* > 0, and select *ϵ*_*D*_, *ϵ*_*T*_> 0 such that *ϵ*_*D*_ + *kϵ*_*T*_ < *ϵ*. By Definition 4, there exists a constant *l*_*D*_ such that NJ applied to matrix *D* computed from sequences of length at least *l*_*D*_ returns *T*\* with probability at least 1 − *ϵ*_*D*_, and there exists a constant *l*_*T*_ such that *M*_*T*_ given DNA sequences of length at least *l*_*T*_ returns *T*\* with probability at least 1 − *ϵ*_*T*_. If a distance matrix *D* is calculated using *M*_*D*_ and a set of *k* constraint trees 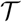 are constructed using *M*_*T*_, given sequences of length at least max{*l*_*D*_, *l*_*T*_}, then the probability that NJ applied to *D* returns *T*\* and that *M*_*T*_ returns a tree that agrees with *T*\* for all *k* constraint trees in 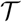 is at least 1 − *ϵ*, as

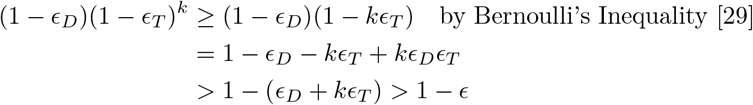

Then, by Theorem 3, NJMerge applied to the input 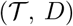 will return the *T*\* with probability at least 1 − *ϵ*, and by Definition 4, NJMerge is statistically consistent under the GTR model.

Finally, we define statistical consistency in the context of species tree estimation (Definition 7) and show that NJMerge can be used to create statistically consistent divide-and-conquer pipelines for species estimation (Corollary 7).

#### Definition 6.

*Let* (*T,* Θ) *be an MSC model tree with topology T and numerical parameters* Θ *(e.g., substitution rate matrix, branch lengths, etc). A method M for constructing species trees from true gene trees is* **statistically consistent** *under the MSC model if, for all ϵ* > 0, *there exists a constant m* > 0 *such that, given at least m true gene trees, M returns T with probability at least* 1 − *ϵ*.

#### Corollary 7.

*NJMerge can be used in a species tree estimation pipeline that is statistically consistent under the MSC model*.

*Proof*. Let (*T*\*, Θ) be an MSC model tree, let *M*_*D*_ be a method for calculating distances between pairs of species from a set of gene trees, and let *M*_*T*_ be a method for constructing species trees from a set of gene trees. Suppose that

- the divide-and-conquer pipeline produces *k* pairwise disjoint subsets of sequences
- Neighbor Joining (NJ) applied to a matrix of pairwise distances calculated using *M*_*D*_ is a statistically consistent method for constructing species trees under the MSC model (e.g., the average topological distance between species in the input set of gene trees [2])
- *M*_*T*_ is statistically consistent under the MSC model (e.g., ASTRAL [27, 28])

Now let *ϵ*> 0, and select *ϵ*_*D*_, *ϵ*_*T*_ > 0 such that *ϵ*_*D*_ + *kϵ*_*T*_ < *ϵ*. By Definition 6, there exists a constant *m*_*D*_ such that NJ applied to matrix *D* computed from at least *m*_*D*_ gene trees returns *T*\* with probability at least 1 − *ϵ*_*D*_, and there exists a constant *m*_*T*_ such that *M*_*T*_ given at least *m*_*T*_ gene trees returns *T*\* with probability at least 1 − *ϵ*_*T*_. If a distance matrix *D* is calculated using *M*_*D*_ and a set of constraint trees 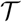 are constructed using *M*_*T*_, both given at least max{*m*_*D*_, *m*_*T*_} gene trees, then the probability that NJ applied to *D* returns *T*\* and that *M*_*T*_ returns a tree that agree with *T*\* for all *k* constraint trees in 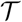 is at least 1 − *ϵ*. Then, by Theorem 3, NJMerge applied to the input 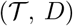 will return the *T*\* with probability at least 1 − *ϵ*, and by Definition 6, NJMerge is statistically consistent under the MSC model.

## Performance Study

Our study evaluated the effectiveness of using NJMerge to estimate species trees on large multi-locus datasets, simulated for this study using the protocol presented in [28]. Our simulation produced model conditions, described by two numbers of taxa (100 and 1000) and two levels of ILS (low/moderate and very high), each with 20 replicate datasets. Datasets included both exon-like sequences and intron-like sequences with exon-like sequences (“exons”) characterized by slower rates of evolution across sites (less phylogenetic signal) and intron-like sequences (“introns”) characterized by faster rates of evolution across sites (greater phylogenetic signal). The 100-taxon datasets were analyzed using 25, 100, and 1000 genes, and the 1000-taxon datasets were analyzed using 1000 genes; note that exons and introns were always analyzed separately. For each of these 320 datasets, we constructed distance matrices using two different methods and constraint trees using four different methods. This provided 2560 different tests on which to evaluate NJMerge. NJMerge failed on 11/2560 tests, so the failure rate (in our experiments) was less than 1%. Species tree methods were evaluated in terms of species tree estimation error (computed using normalized Robinson-Foulds (RF) distances [40]) and running time. All software commands are provided in Additional file 1.

### Simulated Datasets

#### True species and true gene trees

Datasets, each with a true species tree and 2000 true gene trees, were simulated using SimPhy version 1.0.2 [25]. All model conditions had deep speciation (towards the root) and 20 replicate datasets. By holding the effective population size constant (200K) and varying the species tree height (in generations), model conditions with different levels of ILS were generated. For species tree heights of 10M and 500K generations, the average distance between the true species tree and the true gene trees (as measured by the normalized RF distance) was 8–10% and 68–69% respectively. Thus, we referred to these levels of ILS as “low/moderate” and “very high” respectively.

#### True sequence alignments

Sequence alignments were simulated for each true gene tree using INDELible version 1.03 [13] under the GTR+Γ model of evolution without insertions or deletions. For each gene, the parameters for the GTR+Γ model of evolution (base frequencies, substitution rates, and alpha) were drawn from distributions based on estimates of these parameters from the Avian Phylogenomics Dataset [17]; distributions were fitted for exons and introns, separately (Supplementary Table S1, Additional file 1). For each dataset (with 2000 genes), 1000 gene sequences were simulated with parameters drawn from the exon distributions, and 1000 gene sequences were simulated with parameters drawn from the intron distributions. Note that exons and introns were analyzed separately. The sequence lengths were also drawn from a distribution (varying from 300 to 1500 bp).

#### Estimated gene trees

Maximum likelihood gene trees were estimated using FastTree-2 [37] under the GTR+CAT model of evolution. The average gene tree estimation error across all replicate datasets ranged from 26% to 51% for introns and 38% to 64% for exons and thus was higher for exon datasets (Supplementary Table S2, Additional file 1). Note that gene tree estimation error was computed by the normalized symmetric difference between true and estimated gene trees, averaged across all gene trees (the normalized symmetric difference equals the normalized RF distance when both input trees are binary).

### Estimated Species Trees

For each model condition (described by number of taxa and level of ILS), species trees estimation methods were run on the exon-like genes and the intron-like genes, separately. Species trees were estimated on 25, 100, or 1000 genes for the 100-taxon datasets and 1000 genes for the 1000-taxon datasets using three species tree estimation methods: ASTRAL-III [27, 28, 57] (as implemented in version 5.6.1), SVDquartets [9] (as implemented in PAUP* version 4a161 [49]), and concatenation using unpartitioned maximum likelihood under the GTR+Γ model of evolution (as implemented in RAxML [44] version 8.2.12 with pthreads and SSE3).

### NJMerge

#### Distance matrices

Distance matrices were created using two different approaches.

- *D*_*AGID*_ refers to the average gene tree internode distance (AGID) matrix [23], computed from estimated gene trees using ASTRID [51] version 1.1.
- *D*_*LD*_ refers to the log-det distance matrix [46], computed from concatenated alignment using PAUP* [49] version 4a163.

Recall that NJ applied to the AGID matrix (i.e., NJst [23]) was proven to be statistically consistent method under the MSC model [2] and that NJ applied to the log-det distance matrix was proven to be statistically consistent under the MSC model when the sequence evolution models across genes satisfy some additional assumptions (e.g., a relaxed molecular clock) [3].

#### Subset decomposition

We decomposed the species set into subsets as indicated by the blue dashed arrows in Figure 4. Specifically, the NJ tree was computed for each distance matrix using FastME [22] version 2.1.5 and then the centroid tree decomposition (described in PASTA [26]) was used to create disjoint subsets of taxa from the NJ tree. Datasets with 100 species were decomposed into 4–6 subsets with a maximum subset size of 30 taxa, and datasets with 1000 species were decomposed into 10–15 subsets with a maximum subset size of 120 taxa.

#### Constraint trees

Constraint trees were created using four different approaches.

- 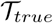 refers to constraint trees computed by restricting the true species tree to each subset of species.
- 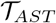 refers to constraint trees computed by running ASTRAL-III on each subset, i.e., on the estimated gene trees restricted to each subset of species.
- 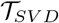 refers to constraint trees computed by running SVDquartets on each subset, i.e., on the concatenated alignment restricted to each subset of species.
- 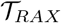 refers to constraint trees computed by running RAxML on each subset, i.e., on the concatenated alignment restricted to each subset of species.

#### Notation

We often specify the inputs to NJ and NJMerge using the following notation: NJ(*D*) and 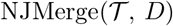. For example, 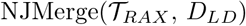 refers to NJMerge given the RAxML constraint trees and the log-det distance matrix as input, whereas 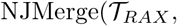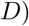 refers to NJMerge given the RAxML constraint trees and **either** the AGID **or** the log-det distance matrix as input.

### Evaluation

#### Species tree estimation error

Species tree estimation error was measured as the RF error rate, i.e., the normalized RF distance between the true and the estimated species trees both on the full species set. Since both trees were fully resolved or binary, the RF error rate is the proportion of edges in the true tree that are missing in the estimated tree. RF error rates were computed using Dendropy [47].

#### Running time

All computational experiments were run on the Blue Waters supercomputer, specifically, the XE6 dual-socket nodes with 64 GB of physical memory and two AMD Interlagos model 6276 CPU processors (i.e., one per socket each with 8 floating-point cores). All methods were given access to 16 threads with 1 thread per bulldozer (floating-point) core. SVDquartets and RAxML were explicitly run with 16 threads; however, ASTRAL-III and NJMerge were not implemented with multi-threading at the time of this study. All methods were restricted to a maximum wall-clock time of 48 hours.

Running time was measured as the wall-clock time and recorded in seconds for all methods. For ASTRAL, SVDquartets, and RAxML, the timing data was recorded for running the method on the full dataset as well as running the method on subsets of the dataset (to produce constraint trees for NJMerge). RAxML did not complete within the maximum wall-clock time of 48 hours on datasets with 1000 taxa, so we used the last checkpoint file to evaluate species tree estimation error and running time. Specifically, running time was measured as the time between the info file being written and the last checkpoint file being written.

We approximated total running time of the NJMerge pipeline by combining the running timing data for estimating the distance matrix, estimating the subset trees, and combining the subset trees using NJMerge. If a user only had access to one compute node, then subset trees would need to be estimated in serial. In this case, the running time of the NJMerge pipeline *t*_*P*_ would be approximated as

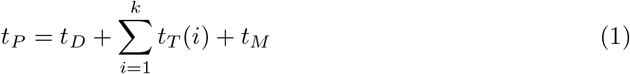

where *k* is the number of subsets, *t*_*D*_ is time to estimate a distance matrix with method *M*_*D*_, *t*_*T*_(*i*) is the time to estimate a species tree on subset *i* with method *M*_*T*_, and *t*_*M*_ is the time to run NJMerge given the distance matrix and the subset trees as input. The average running times for *t*_*T*_ and *t*_*M*_ are shown in Supplementary Tables S9 and S10, Additional file 1. The time to estimate the NJ tree from the distance matrix is not included, as this took less than a minute even for datasets with 1000 species. Note that given access to multiple compute nodes (at least 6 for the 100-taxon datasets and at least 15 for the 1000-species datasets), the subset trees could be estimated in parallel, as shown in [30].

It is worth noting that running ASTRAL-III and computing the AGID matrix requires gene trees to be estimated. Using the same experimental set-up (a single Blue Waters compute node with 64 GB of memory and 16 floating-point cores), FastTree-2 took on average 18 ± 2 minutes to estimate 1000 gene trees for datasets with 100 species and on average 217 ± 20 minutes to estimate 1000 gene trees for datasets with 1000 species (Supplementary Tables S4 and S5, Additional file 1). The amount of time for gene tree estimation can vary greatly, depending on the method used and the analysis performed (e.g., model of sequence evolution, bootstrapping, etc.); we did *not* include the time to estimate gene trees in the reported running times.

## Results

Pipelines using NJMerge can be thought of in two ways: 1) as techniques for potentially improving the accuracy of NJ (hopefully without a large increase in running time) or 2) as techniques for potentially improving the scalability or speed of the method *M*_*T*_ used to compute constraint trees (hopefully without sacrificing accuracy). When distance-based species tree estimation is not as accurate as some other species tree methods, we would predict that NJMerge (when given constraint trees estimated using highly accurate species tree methods) would be more accurate than traditional NJ. Because NJMerge, like NJ, is typically faster than other species tree methods, we would predict that NJMerge would improve the running time of more computationally intensive methods (such as RAxML) used to estimate constraint trees, hopefully without sacrificing accuracy.

Thus, we compared the accuracy of the NJMerge pipeline to traditional NJ, and we also compared the accuracy and running time of the NJMerge pipeline to running *M*_*T*_ on the full dataset, where *M*_*T*_ is the method used to estimate the constraint trees for NJMerge. Results are shown here for intron-like datasets; results for exon-like datasets are shown in Additional file 1. Unless otherwise noted, results were similar for both sequence types; however, species trees estimated on the exon datasets had slightly higher error rates than those estimated on the intron datasets. This is expected, as the exons had slower rates of evolution (and thus less phylogenetic signal) than the introns.

### How do pipelines using NJMerge compare to Neighbor Joining (NJ)?

In this section, we report results on the effectiveness of using NJMerge as compared to NJ in terms of accuracy.

#### Impact of estimated distance matrix

We compared the accuracy of the NJMerge pipeline to traditional NJ on distance matrices estimated from datasets with 100 taxa and varying numbers of genes (Figure 5; Supplementary Figure S1, Additional file 1). Because the accuracy of NJMerge also depends on error in the input constraint trees, we considered an idealized case where NJMerge was given true constraint trees (i.e., constraint trees that agree with the true species tree). We found that 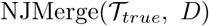 was more accurate than NJ(*D*) for all model conditions and that the difference in error was especially large when the number of genes was small and the level of ILS was very high (e.g., the difference in mean error was greater than 15% when matrices were estimated from 25 introns but was closer to 5% when matrices were estimated from 1000 introns). A similar trend was observed for matrices computed using the log-det distance. Interestingly, both NJ(*D*) and 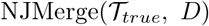 were more accurate when given the AGID matrix rather than the log-det distance matrix as input — even when the level of ILS was low/moderate. In summary, 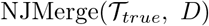 was always more accurate than NJ(*D*), but the improvement in accuracy was greater under challenging model conditions, suggesting that 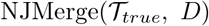 was more robust to error in the distance matrix than NJ(*D*).

**Figure 5:**
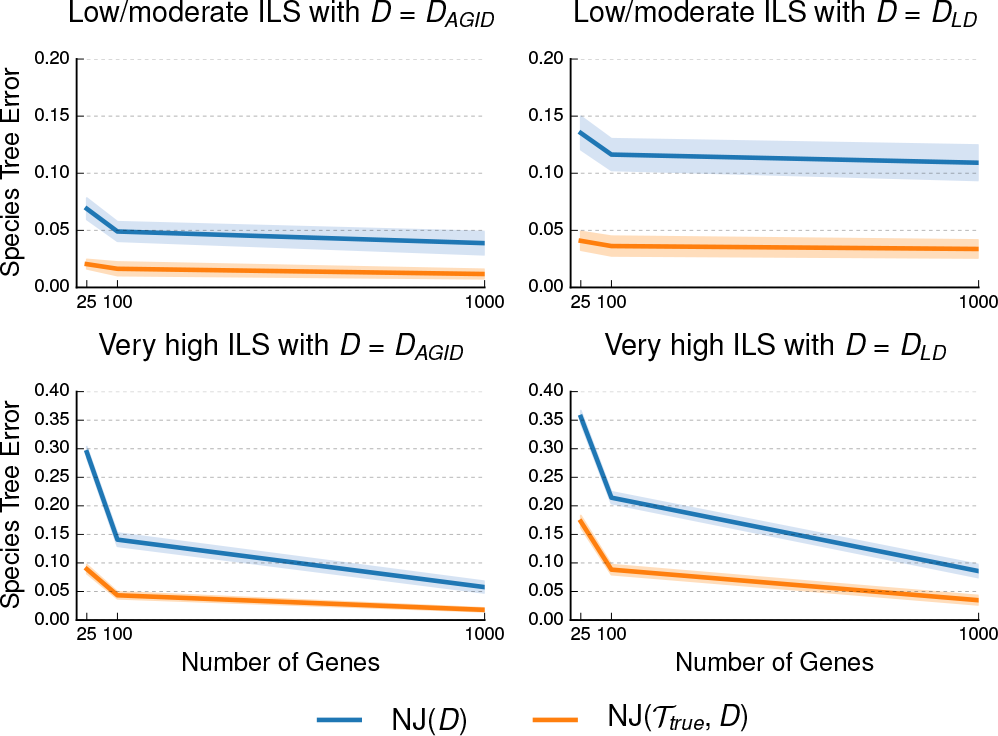
Impact of estimated distance matrix on Neighbor Joining (NJ) and NJMerge. Neighbor Joining (NJ) was run with two different distance matrices, and NJMerge was run with two different distance matrices and constraint trees that agreed with the true species tree (see the Performance Study section for more information on the notation). Datasets had two different levels of incomplete lineage sorting (ILS) and numbers of genes varying from 25 to 1000. Species tree estimation error is defined as the normalized Robinson-Foulds (RF) distance between true and estimated species trees. Lines represent the average over replicate datasets, and filled regions indicate the standard error.

#### Impact of estimated constraint trees

We compared traditional NJ to the NJMerge pipeline given estimated constraint trees on datasets with 1000 taxa and 1000 genes (Figure 6; Supplementary Figure S2, Additional file 1). When the level of ILS was low/moderate, NJMerge outperformed NJ regardless of the method used to estimate species trees. For intron-like datasets with low/moderate ILS, the use of constraint trees reduced the median species tree error from 11–14% (NJ) to less than 3–6% (NJMerge); however, when the level of ILS was very high, the performance of NJMerge varied greatly with the species tree method. Specifically, 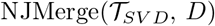 and 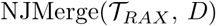 were less accurate than NJ(*D*) by 0-4% on average, whereas 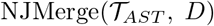 was more accurate than NJ(*D*) by 0–1% on average (Supplementary Tables S7 and S8, Additional file 1). These trends were consistent with the relative performance of methods on the 100-taxon datasets (Figure 7 and Supplementary Figure S3, Additional file 1); specifically, when the level of ILS was very high, SVDquartets and RAxML performed worse than running NJ on *either* the AGID matrix or the log-det distance matrix. In summary, NJMerge was highly impacted by the quality of the constraint trees — so that accurate constraint trees resulted in NJMerge being more accurate than NJ, but inaccurate constraint trees resulted in NJMerge being less accurate than NJ.

**Figure 6:**
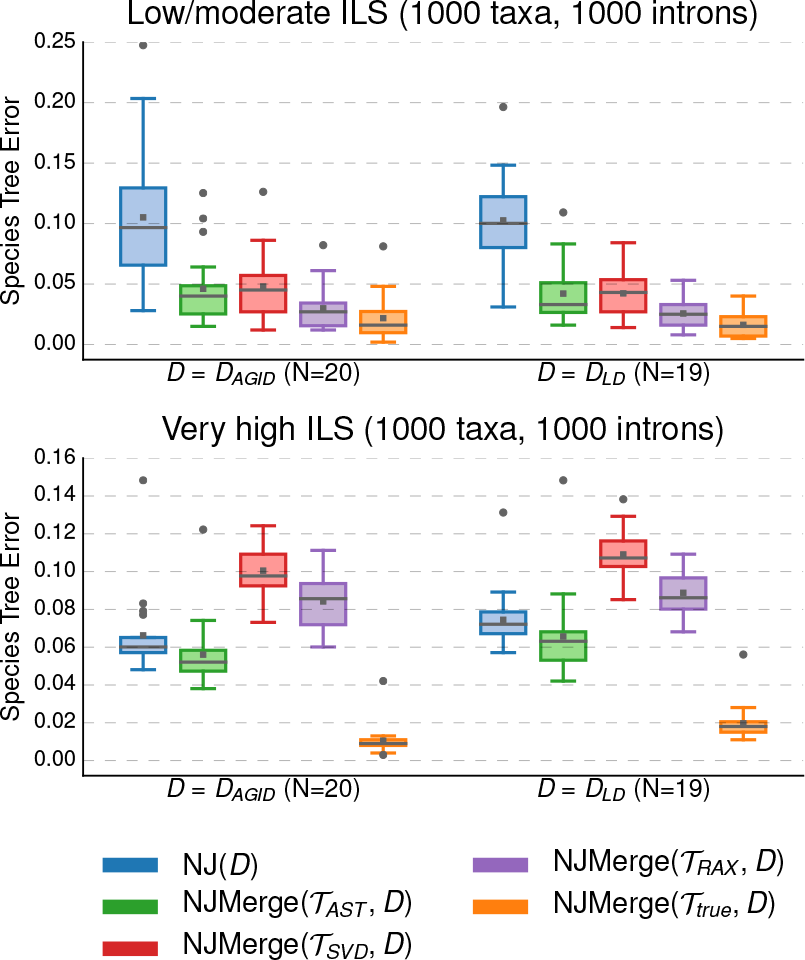
Impact of estimated constraint trees on NJMerge. Neighbor Joining (NJ) was run with two different distance matrices, and NJMerge was run with two different distance matrices and four different sets of constraint trees (see the Performance Study section for more information on the notation). Species tree estimation error is defined as the normalized Robinson-Foulds (RF) distance between true and estimated species trees. Note that gray bars represent medians, gray squares represent means, gray circles represent outliers, box plots are defined by quartiles (extending from the first to the third quartiles), and whiskers extend to plus/minus 1.5 times the interquartile distance (unless greater/less than the maximum/minimum value).

**Figure 7:**
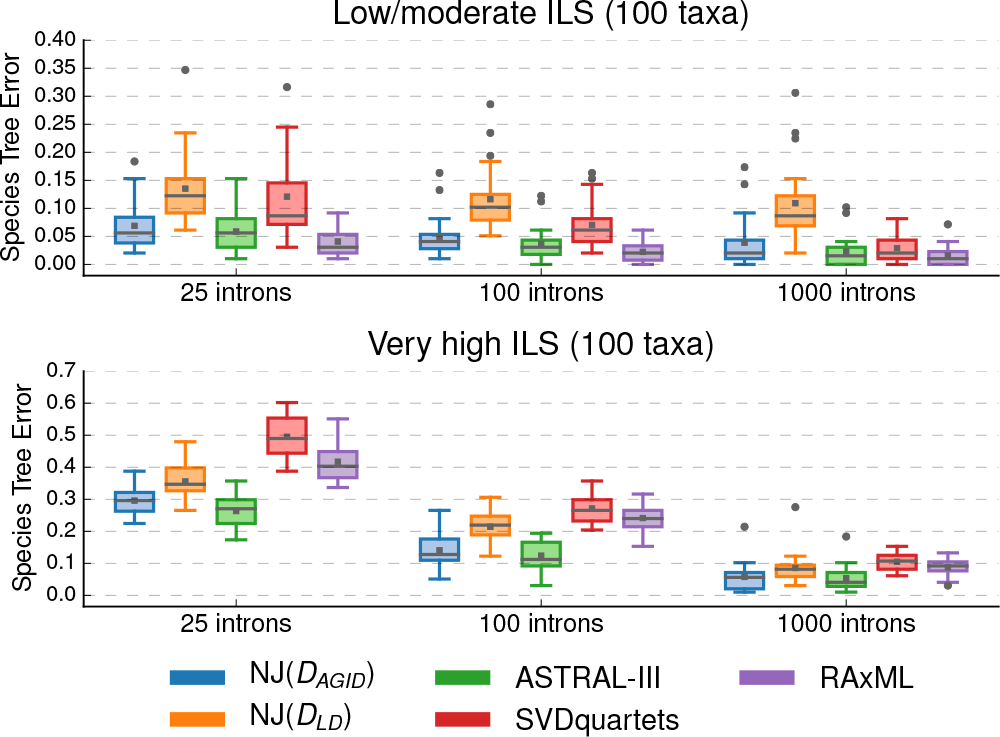
Comparison of species tree methods. All methods were run on the full dataset (i.e., not subsets) with 100 species. Neighbor Joining (NJ) was run with two different distance matrices (Performance Study section for more information on the notation). Species tree estimation error is defined as the normalized Robinson-Foulds (RF) distance between true and estimated species trees. Note that gray bars represent medians, gray squares represent means, gray circles represent outliers, box plots are defined by quartiles (extending from the first to the third quartiles), and whiskers extend to plus/minus 1.5 times the interquartile distance (unless greater/less than the maximum/minimum value).

### How do pipelines using NJMerge compare to ASTRAL-III, SVDquartets, and RAxML?

In this section, we compare the running time and the accuracy of the NJMerge pipeline to running *M*_*T*_ on the full dataset, where *M*_*T*_ is the method used to estimate constraint trees for NJMerge. Because NJMerge was more accurate when given the AGID matrix (Figure 5; Supplementary Figure S1, Additional file 1), results for NJMerge given the AGID distance matrix are shown here, and results for NJMerge given the log-det distance matrix are shown in Additional file 1.

#### ASTRAL-III vs. NJMerge

Both 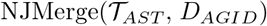 and 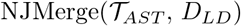 provided running time advantages over ASTRAL-III under some model conditions. While ASTRAL-III completed on all the low/moderate ILS datasets with 1000 taxa and 1000 genes in less than 9 hours on average, ASTRAL-III failed to complete within the maximum wall-clock time of 48 hours on 23/40 datasets with 1000 taxa, 1000 genes, and very high ILS (Table 1). On the other 17/40 datasets, ASTRAL-III ran for more than 2000 minutes (approximately 33 hours). This difference between the low/moderate ILS and the very high ILS datasets is noteworthy (see discussion). In contrast, 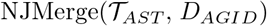 completed in under 300 minutes (approximately 5 hours) on average, including the time it took to estimate the distance matrix and the ASTRAL-III subset trees in serial (Figure 8, Supplementary Figure S4, Additional file 1). Note that 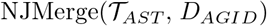 failed on 0 datasets, and 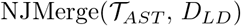 failed on 2 datasets (Table 1). In summary, NJMerge substantially reduced the running time of ASTRAL-III on the 1000-taxon, 1000-gene datasets with very high ILS.

**Table 1:** The number of datasets on which methods failed is indicated below by model condition. ASTRAL-III failed due to running beyond the maximum wall clock time of 48 hours; SVDquartets failed due to segmentation faults; RAxML failed due to running out of memory; NJMerge failed due to being unable to find a legal siblinghood. Note that NJMerge is described by the input set 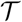 of constraint trees and input distance matrix *D*; see the Performance Study section for more information on the notation.

| # of<br>Taxa | # of<br>Genes | ILS<br>Level | Sequence<br>Type | Method | # of Failures<br>(out of 20) |
| --- | --- | --- | --- | --- | --- |
| 100 | 25 | very high | exon | NJMerge($\mathcal{T}_{true}, D_{LD}$) | 1 |
| 100 | 25 | very high | exon | NJMerge($\mathcal{T}_{RAX}, D_{AGID}$) | 1 |
| 100 | 25 | very high | intron | NJMerge($\mathcal{T}_{true}, D_{AGID}$) | 1 |
| 1000 | 1000 | low/moderate | exon | SVDquartets | 20 |
| 1000 | 1000 | low/moderate | exon | RAxML | 3 |
| 1000 | 1000 | low/moderate | intron | NJMerge($\mathcal{T}_{AST}, D_{LD}$) | 1 |
| 1000 | 1000 | low/moderate | intron | SVDquartets | 20 |
| 1000 | 1000 | low/moderate | intron | RAxML | 20 |
| 1000 | 1000 | very high | exon | ASTRAL-III | 19 |
| 1000 | 1000 | very high | exon | NJMerge($\mathcal{T}_{true}, D_{LD}$) | 1 |
| 1000 | 1000 | very high | exon | NJMerge($\mathcal{T}_{AST}, D_{LD}$) | 1 |
| 1000 | 1000 | very high | exon | NJMerge($\mathcal{T}_{SVD}, D_{LD}$) | 2 |
| 1000 | 1000 | very high | exon | NJMerge($\mathcal{T}_{RAX}, D_{LD}$) | 2 |
| 1000 | 1000 | very high | exon | SVDquartets | 20 |
| 1000 | 1000 | very high | intron | ASTRAL-III | 4 |
| 1000 | 1000 | very high | intron | NJMerge($\mathcal{T}_{SVD}, D_{LD}$) | 1 |
| 1000 | 1000 | very high | intron | SVDquartets | 20 |
| 1000 | 1000 | very high | intron | RAxML | 19 |

**Figure 8:**
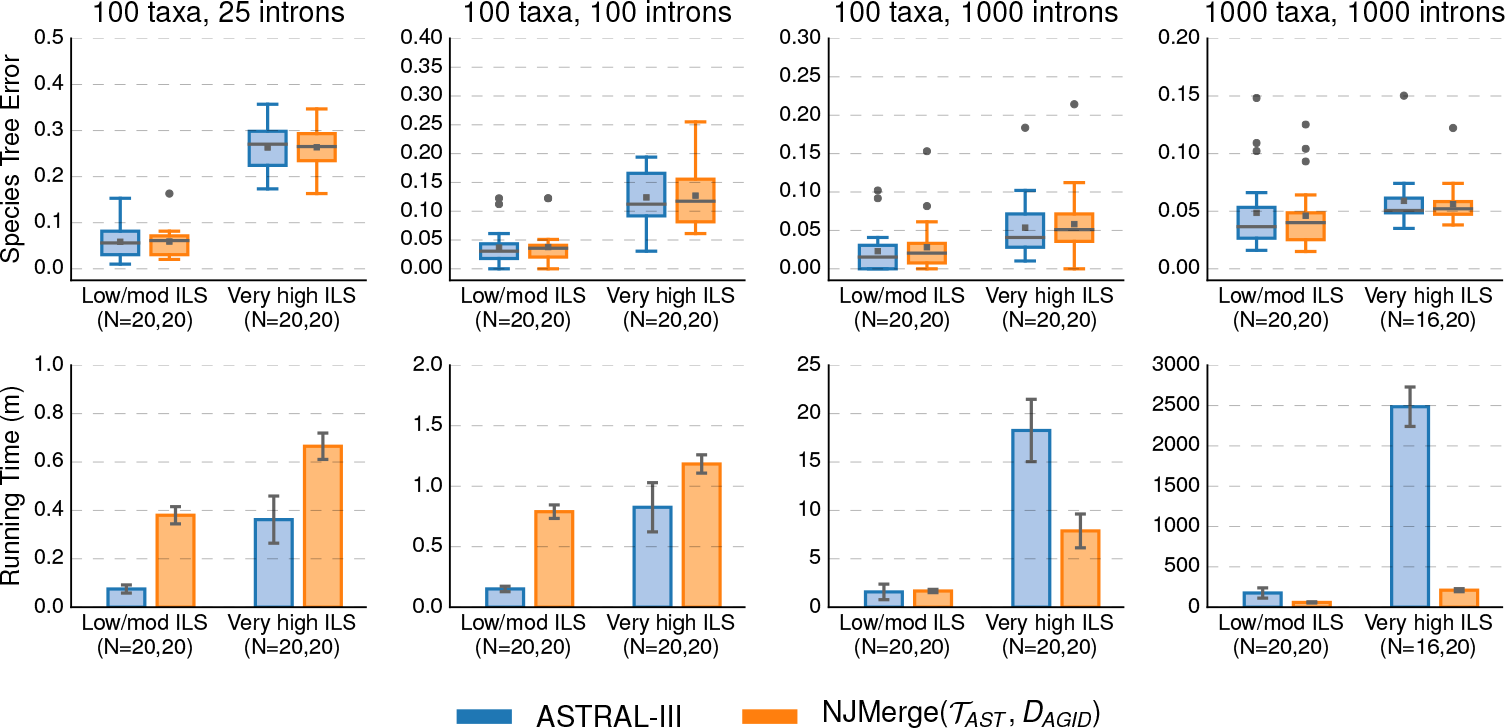
ASTRAL-III vs. NJMerge given ASTRAL-III constraint trees and average gene tree internode distance (AGID) matrix. Subplots on top row show species tree estimation error (defined as the normalized RF distance between true and estimated species trees); note that gray bars represent medians, gray squares represent means, gray circles represent outliers, box plots are defined by quartiles (extending from the first to the third quartiles), and whiskers extend to plus/minus 1.5 times the interquartile distance (unless greater/less than the maximum/minimum value). Subplots on bottom row show running time (in minutes); bars represent means and error bars represent standard deviations across replicate datasets. NJMerge running times is for computing the subset trees “in serial”; see Equation (1) in the main text for more information. The numbers of replicates on which the methods completed is shown on the x-axis, e.g., *N* = *X, Y* indicates that ASTRAL-III completed on *X* out of 20 replicates and that 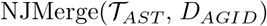 completed on *Y* out of 20 replicates. ASTRAL-III did not complete within the maximum wall-clock time of 48 hours on 4/40 intron-like datasets with 1000 taxa and very high ILS.

ASTRAL-III and 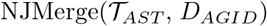 achieved similar levels of accuracy with the mean species tree error within 0–2% for both intron and exon datasets (Figure 8; Supplementary Figure S4, Additional file 1; Supplementary Table S7, Additional file 1). Trends were similar for 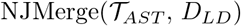 except when the level of ILS was very high; under these conditions, the mean error of 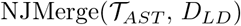 was 2–6% greater than that of ASTRAL-III (Supplementary Figures S7 and S8, Additional file 1; Supplementary Table S8, Additional file 1).

#### NJMerge vs. SVDquartets

Species trees can be estimated with SVDquartets using the full set of 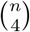 quartet trees or a subset of quartet trees. Based on a prior study [48], which showed that the best accuracy was obtained when using all quartet trees, we computed all 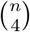 quartet trees for 100-taxon datasets. However, on datasets with 1000 taxa, SVDquartets was run using a random subset of quartet trees (without replacement), because the maximum number of quartets allowed by SVDquartets (as implemented by PAUP*) was 4.15833 × 10^10^. Running PAUP* resulted in a segmentation fault for all 1000-taxon datasets, i.e., SVDquartets failed on 40/40 datasets with 1000 taxa and 1000 genes.

In contrast, 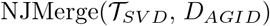 failed on 0 datasets, and 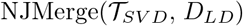 failed on 3 datasets (Table 1).

NJMerge also improved running time on datasets with 100 taxa; for example, SVDquartets completed in 19–81 minutes on average, whereas 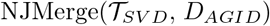 completed in less than 2 minutes on average for datasets with 100 taxa and 1000 genes (Figure 9; Supplementary Figure S5, Additional file 1). This running time comparison does not take into account the time needed to estimate gene trees, which required on average 18 minutes using FastTree-2 on datasets with 100 taxa and 1000 genes.

**Figure 9:**
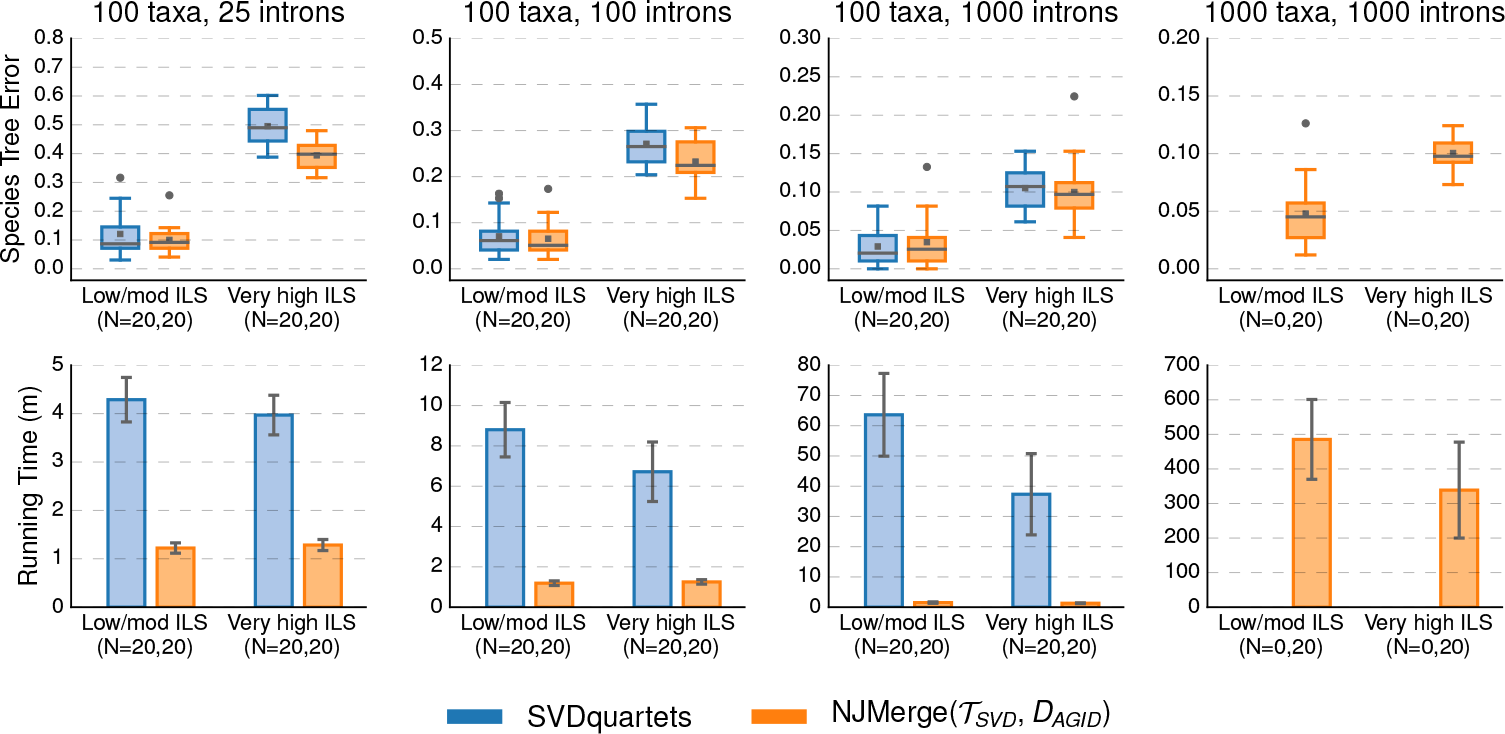
SVDquartets vs. NJMerge given SVDquartet constraint trees and average gene tree internode distance (AGID) matrix. Subplots on top row show species tree estimation error (defined as the normalized RF distance between true and estimated species trees); note that gray bars represent medians, gray squares represent means, gray circles represent outliers, box plots are defined by quartiles (extending from the first to the third quartiles), and whiskers extend to plus/minus 1.5 times the interquartile distance (unless greater/less than the maximum/minimum value). Subplots on bottom row show running time (in minutes); bars represent means and error bars represent standard deviations across replicate datasets. NJMerge running times is for computing the subset trees “in serial”; see Equation (1) in the main text for more information. The numbers of replicates on which the methods completed is shown on the x-axis, e.g., *N* = *X, Y* indicates that SVDquartets completed on *X* out of 20 replicates and that 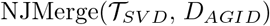 completed on *Y* out of 20 replicates. SVDquartets did not run any datasets with 1000 taxa due to segmentation faults.

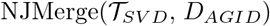 typically produced species trees with less error than SVDquartets. The difference between methods was typically small (between 0–2%) when the level of ILS was low/moderate but could be larger than 10% when the level of ILS was very high. Similar trends were observed for 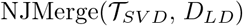 (Supplementary Figures S9 and S10, Additional file 1).

#### NJMerge vs. RAxML

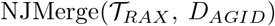 and 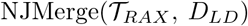 reduced the running time of RAxML by more than half — even though RAxML was run on the subset trees in serial (Figure 10 and Supplementary Figure S6, Additional file 1). For the 1000-taxon datasets, the final checkpoint was written by RAxML after more than 2250 minutes (~37.5 hours) on average. In comparison, when RAxML was run on subsets in serial, the average running time of 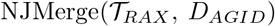 was between 500 (approximately 8.5 hours) and 1500 minutes (approximately 25 hours). Although these running times for NJMerge do not include the time to estimate gene trees, recall that it took on average 217 minutes (less than 4 hours) to estimate 1000 gene trees on datasets with 1000 species using FastTree-2.

**Figure 10:**
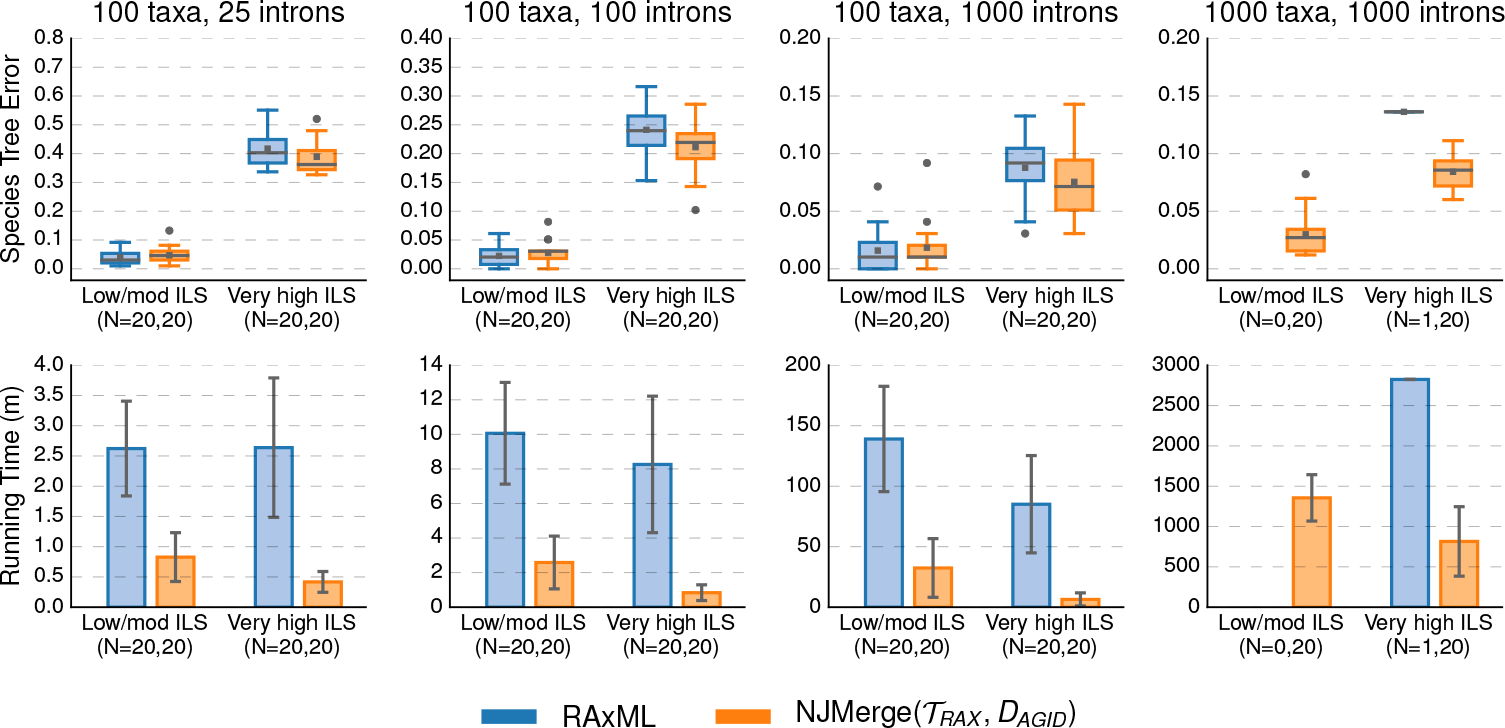
RAxML vs. NJMerge given RAxML constraint trees and and average gene tree internode distance (AGID) matrix. Subplots on top row show species tree estimation error (defined as the normalized RF distance between true and estimated species trees); note that gray bars represent medians, gray squares represent means, gray circles represent outliers, box plots are defined by quartiles (extending from the first to the third quartiles), and whiskers extend to plus/minus 1.5 times the interquartile distance (unless greater/less than the maximum/minimum value). Subplots on bottom row show running time (in minutes); bars represent means and error bars represent standard deviations across replicate datasets. NJMerge running times is for computing the subset trees “in serial”; see Equation (1) in the main text for more information. The numbers of replicates on which the methods completed is shown on the x-axis, e.g., *N* = *X, Y* indicates that RAxML completed on *X* out of 20 replicates and that 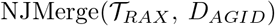 completed on *Y* out of 20 replicates. RAxML was only able to run on 1/40 intron-like datasets with 1000 taxa due to “Out of Memory” errors.

While NJMerge can fail to return a tree, NJMerge failed less frequently than RAxML — when both methods were given the same computational resources. 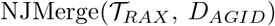 failed on 1 dataset, and 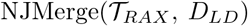 failed on 2 datasets. In contrast, for datasets with 1000 taxa, RAxML failed to run on 38 intron-like datasets and 3 exon-like datasets due to “Out of Memory” (OOM) errors (Table 1); the difference between the number of intron-like versus the number of exon-like datasets is noteworthy (see discussion).

For datasets with low/moderate levels of ILS, RAxML produced species trees with less error (0–3% on average) than 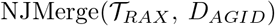; however, for datasets with very high levels of ILS, 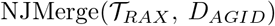 produced species trees with less error 0–4% on average) than RAxML (Figure 10; Supplementary Figure S3, Additional file 1). Similar trends were observed for 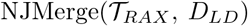 (Supplementary Figures S11 and S12, Additional file 1).

## Discussion

### Remarks on the utility of pipelines using NJMerge

Pipelines using NJMerge can be viewed either as techniques for improving traditional NJ or as techniques for scaling a computationally-intensive base method (previously referred to as *M*_*T*_) to larger datasets. Thus, in order to maximize the utility of NJMerge, users should select a base method that is both more accurate and more computationally-intensive than NJ. Our results show that selecting base methods for NJMerge may not be trivial when analyzing phylogenomic datasets — because both accuracy and running time were impacted by the level of ILS. For example, ASTRAL-III was very fast when the level of ILS was low/moderate but was substantially slower when the level of ILS was very high. Similarly, SVDquartets and RAxML were both more accurate than NJ(*D*_*AGID*_), i.e., NJst, when the level of ILS was low/moderate but were less accurate than these methods when the level of ILS was very high; note that this trend is consistent with results from [31] (also see the review paper by [56]). Overall, our results suggest that constraint trees should be estimated using RAxML when the level of ILS is low/moderate and using ASTRALIII when the level of ILS is very high, and thus, determining whether the level of ILS in a given phylogenomic datasets is an important area of future research. Finally, we note that NJMerge, when given constraint trees that agreed with the true species tree, was very accurate (less than 2% error on average) even when the level of ILS was very high, suggesting that NJMerge is a promising technique for scaling Bayesian methods (e.g., Starbeast2 [35]) and future species tree methods to larger datasets.

Although NJMerge can fail, this should not discourage potential users, as NJMerge failed on fewer datasets than ASTRAL-III, SVDquartets, or RAxML — when all methods were given the same computational resources, including a maximum wall-clock time of 48 hours. In our experiments, NJMerge failed on only 11/2560 test cases from running NJMerge on 320 datasets with two different types of distance matrices and four different types of constraint trees (Table 1).

Importantly, in all our experiments, NJMerge was run within the divide-and-conquer pipeline shown in Figure 4, specifically, with subsets of taxa derived from decomposing the NJ tree (blue dashed lines). Because NJMerge was always given inputs generated by this pipeline, our results on the accuracy, the failure rate, and the running time of NJMerge may not generalize to arbitrary inputs.

### Remarks on other results

#### Impact of distance matrix on NJ

Our results showed that on average NJ(*D*_*AGID*_) was either as accurate or else more accurate than NJ(*D*_*LD*_). Notably, there was a clear difference between these two methods on datasets with 100 taxa and low/moderate levels of ILS; specifically NJ(*D*_*AGID*_) produced trees with less than 5% error on average, whereas NJ(*D*_*LD*_) produced trees with greater than 10% error on average). However, on the exact same model condition but with 1000 taxa, NJ(*D*_*AGID*_) and NJ(*D*_*LD*_) produced trees with similar levels of accuracy. This may be due to the difference between the median branch length between low/moderate ILS datasets with 100 taxa and 1000 taxa (Supplementary Table S3, Additional file 1); furthermore, it is possible that branch length and other factors that limit the accuracy of NJ(*D*_*LD*_) in the context of gene tree estimation would also apply in the context of species tree estimation. However, it is interesting to note that NJ(*D*_*LD*_) was more accurate than either SVDquartets or RAxML when the level of ILS was very high, providing support for Allman et al.’s statement, “The simplicity and speed of distancebased inference suggests log-det based methods should serve as benchmarks for judging more elaborate and computationally-intensive species trees inference methods” [3].

#### Impact of ILS and sequence type on ASTRAL-III

Our results showed that ASTRAL-III was much faster on the low/moderate ILS datasets than on the very high ILS datasets. This finding makes sense in light of ASTRAL-III’s algorithm design. ASTRAL-III operates by searching for an optimal solution to its search problem within a constrained search space that is defined by the set 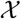 of bipartitions in the estimated gene trees, and in particular, ASTRAL-III’s running time scales with 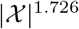 [57]. The set of gene trees will become more heterogeneous for higher levels of ILS, and thus, the size of 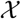 will increase, as every gene tree could be different when the level of ILS is very high. In addition, gene tree estimation error can also increase the size of 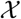, explaining why ASTRAL-III failed to complete on exon datasets more often than on intron datasets (Table 1 Supplementary Table S2, Additional file 1).

#### Impact of sequence type on RAxML

Our results showed that RAxML failed on more intron-like datasets than exon-like datasets. This finding makes sense in light of RAxML’s implementation. RAxML uses redundancy in site patterns to store the input alignment compactly, so that the memory scales with the number of unique site patterns. The intron datasets had more unique site patterns than the exon datasets (i.e., greater phylogenetic signal and lower gene tree estimation error), which explains why RAxML required more memory when analyzing introns.

### Remarks on the statistical consistency of pipelines using NJMerge

Although NJMerge can fail to return a tree, by statistical consistency under the MSC model (Corollary 7), the probability that NJMerge fails goes to zero as the number of true gene trees goes to infinity. In fact, NJMerge was designed to have this theoretical guarantee via the selection of the heuristic for determining whether or not to accept a siblinghood proposal. It is easy to think of other heuristics that prevent NJMerge from failing but do *not* have the guarantee of correctness (Theorem 3) and thus do *not* have the guarantee of statistical consistency (Corollary 7). Designing heuristics that prevent NJMerge from failing but have good theoretical properties is an area of future research.

As mentioned previously, our proof of statistical consistency under the MSC model requires that the number of true gene trees goes to infinity, which is the equivalent of requiring that *both* the number of gene trees and the sequence length per gene tree go to infinity. Roch et al. [41] recently showed that essentially all gene tree summary methods (e.g., NJst [2], and ASTRAL [27]) are not statistically consistent under the MSC if the sequence length per gene is fixed — and these theoretical results apply to NJMerge as well. The failure to be statistically consistent when the sequence length per gene is bounded is not unique to gene tree summary methods or NJMerge, as Roch et al. also showed that fully partitioned maximum likelihood is not consistent under these conditions, and [42] had shown that unpartitioned maximum likelihood is also not consistent.

## Conclusions

In this paper, we introduced a divide-and-conquer approach to phylogeny estimation that 1) decomposes a set of species into pairwise disjoint subsets, 2) builds trees on each subset of species using a base method, and 3) merges the subsets trees together using a distance matrix. For the merger step, we presented a new method, called NJMerge, and proved that some divide-and-conquer pipelines using NJMerge are statistically consistent under some models of evolution. We then evaluated pipelines using NJMerge in the context of species tree estimation, specifically using simulated multi-locus datasets with up to 1000 species and two levels of ILS. We found that pipelines using NJMerge provided several benefits to large-scale species tree estimation. Specifically, under some model conditions, pipelines using NJMerge improved the accuracy of traditional NJ and substantially reduced the running time of three popular species tree methods (ASTRAL-III, SVDquartets, and “concatenation” using RAxML) without sacrificing accuracy (see discussion for details as the results depended on the level of ILS). Finally, although NJMerge can fail to return a tree, in our experiments, pipelines using NJMerge failed on only 11 out of 2560 test cases. Together these results suggest that NJMerge is a promising approach for scaling highly accurate but computationally-intensive methods to larger datasets.

This study also suggests several different directions for future research. Since NJMerge uses a heuristic (which can fail) to test for tree compatibility (in deciding whether to accept a siblinghood proposal), a modification to NJMerge to use an exact method for this problem would reduce the failure rate and — if sufficiently fast — would still enable scalability to large datasets. In addition, all aspects of the divide-and-conquer pipeline could be modified and tested; for example, the robustness of NJMerge to the starting tree and initial subset decomposition could be evaluated. Finally, divide-and-conquer pipelines using NJMerge could be compared to traditional divide-and-conquer pipelines (e.g., Disk Covering Methods) when robust implementations become publicly available for species tree estimation. Other agglomerative techniques for merging disjoint subset trees are being developed (e.g., the agglomerative technique described in [58] for gene tree estimation has good theoretical properties but has not yet been implemented), and NJMerge should be compared to such techniques when they become publicly available.

## Abbreviations

*GTR*: Generalized Time Reversible
*ILS*: Incomplete Lineage Sorting
*MSC*: Multi-Species Coalescent
*NJ*: Neighbor Joining
*RF*: Robinson-Foulds

## Availability and requirements

Project name: NJMerge; Project home page: https://github.com/ekmolloy/njmerge; Operating systems: Platform independent; Programming language: Python version 2.7; Other requirements: Dendropy version 4.3.0; License: BSD 3-Clause; Any restrictions to use by non-academics: None.

## Availability of data and materials

The datasets supporting the conclusions of this article are available in the following Illinois Data Bank repositories: https://doi.org/10.13012/B2IDB-7735354_V1 and https://doi.org/10.13012/B2IDB-0569467_V1.

## Ethics approval and consent to participate

Not applicable.

## Consent for publication

Not applicable.

## Author’s contributions

Both authors designed the study, proved the theorems, and wrote the paper. EKM implemented the algorithm and performed the simulation study.

## Acknowledgements

We thank the anonymous reviewers of our RECOMB-CG paper, whose feedback led to improvements in the quality of this paper.

