## Supplementary Materials for "Statistically consistent divide-and-conquer pipelines for phylogeny estimation using NJMerge"

Erin K. Molloy and Tandy Warnow

April 20, 2019

#### Contents

**List of Tables** 1

**List of Figures** 1

**References** 26

S9 Comparison of SVDquartets and NJMerge given log-det distance matrix for intron-like datasets 18
S10 Comparison of SVDquartets and NJMerge given log-det distance matrix for exon-like datasets 19
S11 Comparison of RAxML and NJMerge given log-det distance matrix for intron-like datasets . 20
S12 Comparison of RAxML and NJMerge given log-det distance matrix for exon-like datasets . . 21

### Supplementary Methods

#### Simulation Command

SimPhy [7] Version 1.0.2 was run as

```
simphy -rs 3000 -rl F:3000 -rg 1 -st F:[species tree height] \
       -si F:1 -sl F:[number of taxa] -sb F:0.0000001 -sp F:200000 \
       -hs LN:1.5,1 -hl LN:1.2,1 -hg LN:1.4,1 -su E:10000000 \
       -so F:1 -od 1 -v 3  -cs 293745 -o [output directory name]
```

where the number of taxa was 100 or 1000, the species tree height was 10,000,000 or 500,000 generations, and the effective population size was constant at 200,000.

#### INDELible Simulation

Like the simulation in [8], GTR+$\Gamma$ model parameters (base frequencies, substitution rates, and alpha) were drawn from distributions. However, unlike the simulation in [8], we estimated separate distributions for exons, introns, and ultra-conserved elements (UCEs) from the Avian Phylogenomics Dataset [4]. GTR model parameters that were drawn from distributions given in Table S1. The $\alpha$ parameter for the $\Gamma$ distribution was computed as follows. (1) For each gene, a random value $x$ between 0 and 1 is drawn from a uniform distribution. (2) $\alpha$ was computed as $-log(x)/d$, where $d$ is 4.2 for exons, 0.4 for introns, and 1.0 for UCEs. Because the mean of the uniform distribution is 0.5, the mean $\alpha$ is approximately 0.16 for exons, 1.73 for introns, and 0.69 for UCEs. We ran INDELible [3] Version 1.03 using custom Python scripts available on the Illinois Data Bank using the exon, intron, and UCE parameter distributions for genes 1-1000, 1001-2000, and 2001-3000, respectively. Only the exon-like and intron-like genes were used in this study due to limitations in computational resources.

Table S1: **INDELible Simulation Parameters.**

| Sequence Type | GTR Base Frequencies | GTR Substitution Rates |
|---|---|---|
| Exon | Dirchlet(79,57,60,53) | Dirchlet(3,9,2,4,11,7) |
| Intron | Dirchlet(55,38,43,63) | Dirchlet(37,133,20,48,120,29) |
| UCE | Dirchlet(68,45,45,68) | Dirchlet(19,66,10,27,67,19) |

#### Average Gene-Tree Internode Distance Matrix Commands

FastTree [10] Version 2.1.10 (SSE3) was run as

```
FastTree -nt -gtr -quiet -log fasttree-$gene.log \
     [input alignment fasta file] > [output FastTree-2 tree file]
```

ASTRID [15] Version 1.4 was run as

```
ASTRID -i [input gene tree list file] \
       -c [output distance matrix file] -o [temporary file]
```

#### LogDet Distance Matrix Command

PAUP* [14] 4a163 64-bit Centos was run as

```
echo  "ToNEXUS format=FASTA fromFile=[input alignment fasta file]
    toFile=[alignment nexus file]; exe [alignment nexus file]; DSet distance=logdet;
    SaveDist format=PHYLIP file=[output distance matrix file] triangle=both diagonal=yes;" |
    paup4a163_centos64 -n
```

#### Species Tree Commands

ASTRAL [16] Version 5.6.1 (i.e., ASTRAL-III) was run as

```
java java -Xms3200M -Xmx32000M ASTRAL/Astral/astral.5.6.1.jar \
     -i [input gene tree list file] -o [output ASTRAL-III tree file]
```

SVDquartets [1, 2] (PAUP* [14] Version 4a161/3) was run as

```
echo "exe [input alignment nexus file]; svd nthreads=16
    evalQuartets=all qfile=[output quartet file] qformat=qmc;
    savetrees file=[output SVDquartets tree file] format=newick;" |
    paup4a161_centos64 -n
```

PAUP* [14] Version 4a161 64-bit Centos was used for results obtained in [9], and PAUP* Version 4a163 64-bit Centos was used for results obtained using log-det distance matrix.

RAxML [11] Version 8.2.12 (with pThreads SSE3) was run as

```
raxmlHPC-PTHREADS-SSE3 -m GTRGAMMA -F -p [seed] \
    -n [output name] -s [input alignment file] -T 16
```

Note that the option `-j` (to write checkpoints) was included for the 1000-taxon datasets only.

#### NJ / NJMerge Commands

Neighbor Joining (FastME [5] Version 2.1.5) was run as

```
fastme -mN -i [input distance matrix file] -o [output tree file]
```

NJMerge was run as

```
python njmerge.py \
    -t [input constraint tree file 1] ... [input constraint tree file N] \
    -m [input internode distance matrix file] \
    -x [input rows to taxon name map file] \
    -o [output NJMerge tree file]
```

#### Tree Comparison Commands

The normalized symmetric difference was computed using Dendropy [13] Version 4.3.0 as

```
n1 = len(t1.internal_edges(exclude_seed_edge=True))
n2 = len(t2.internal_edges(exclude_seed_edge=True))
[fp, fn] = false_positives_and_negatives(t1, t2)
sd = float(fp + fn) / (n1 + n2)
```

where `t1` and `t2` are Dendropy tree objects. Note that the normalized symmetric difference equals the normalized Robinson-Foulds distance when both trees are fully resolved (i.e., binary). In this study, both true and estimated species trees as well as true gene trees were fully resolved; however, gene trees estimated using FastTree-2 could have polytomies.

#### Simulated Datasets

Table S2: **Simulated Dataset Properties.** Simulated datasets are described by the average discord between the species tree and the gene trees as well as the gene tree estimation error. Specifically, "Average Distance" is the normalized symmetric difference distance between the true species tree and the true gene tree, averaged across all 1000 genes in a replicate dataset. "Gene Tree Estimation Error" is the normalized symmetric difference between the true and the estimated gene trees, averaged across all 1000 genes in a replicate dataset. "Total Gene Tree Discord" is the normalized symmetric difference between the true species tree and the estimated gene tree, averaged across all 1000 genes in a replicate dataset. Values below are the mean (± standard deviation) across 20 replicates.

| Number of Taxa | Sequence Type | Average Distance | Gene Tree Estimation Error | Total Gene Tree Discord |
|---|---|---|---|---|
| *Moderate ILS (species tree height = 10M generations)* | | | | |
| 100 | exon | $0.08 \pm 0.02$ | $0.38 \pm 0.06$ | $0.39 \pm 0.06$ |
| 100 | intron | $0.08 \pm 0.02$ | $0.26 \pm 0.07$ | $0.28 \pm 0.06$ |
| 1000 | exon | $0.10 \pm 0.00$ | $0.42 \pm 0.04$ | $0.43 \pm 0.04$ |
| 1000 | intron | $0.10 \pm 0.00$ | $0.30 \pm 0.05$ | $0.32 \pm 0.05$ |
| *Very High ILS (species tree height = 500K generations)* | | | | |
| 100 | exon | $0.68 \pm 0.02$ | $0.57 \pm 0.07$ | $0.78 \pm 0.03$ |
| 100 | intron | $0.68 \pm 0.02$ | $0.43 \pm 0.10$ | $0.74 \pm 0.03$ |
| 1000 | exon | $0.69 \pm 0.01$ | $0.64 \pm 0.05$ | $0.81 \pm 0.02$ |
| 1000 | intron | $0.69 \pm 0.01$ | $0.51 \pm 0.07$ | $0.76 \pm 0.03$ |

Table S3: **Gene Tree Median Branch Lengths.** For the first replicate dataset in each model condition, we computed the median branch length per gene tree for the internal branches as well as the terminal branches, separately. Values below are the mean (± standard deviation) across 1000 gene trees.

| Number of Taxa | Sequence Type | Internal Branch Lengths | Terminal Branch Lengths |
|---|---|---|---|
| *Moderate ILS (species tree height = 10M generations)* | | | |
| 100 | exon | $0.0317 \pm 0.0235$ | $0.0588 \pm 0.0436$ |
| 100 | intron | $0.0317 \pm 0.0229$ | $0.0585 \pm 0.0425$ |
| 1000 | exon | $0.0255 \pm 0.0184$ | $0.0527 \pm 0.0378$ |
| 1000 | intron | $0.0260 \pm 0.0189$ | $0.0535 \pm 0.0389$ |
| *Very High ILS (species tree height = 500K generations)* | | | |
| 100 | exon | $0.0012 \pm 0.0009$ | $0.0072 \pm 0.0051$ |
| 100 | intron | $0.0012 \pm 0.0010$ | $0.0073 \pm 0.0055$ |
| 1000 | exon | $0.0010 \pm 0.0007$ | $0.0073 \pm 0.0053$ |
| 1000 | intron | $0.0009 \pm 0.0007$ | $0.0066 \pm 0.0048$ |

#### Approximation of Running Time for Gene Tree Estimation

We *re-estimated* gene trees in order to include the average time required for gene tree estimation for each replicate dataset (either 1000 exons or 1000 introns). Jobs were submitted using the qsub script below, where `$modl` is the model condition (e.g., `100tax-3000gen-10M`) and `$repl` is the replicate number (e.g., `08`). Technically, the script below estimates gene trees on gene numbered 1 to 1000 (i.e., 1000 exons), and a similar script was used to estimate gene trees numbered 1001 to 2000 (i.e., 1000 introns). This script will "launch serial tasks, up to [16] running concurently on a single node. Any number of tasks can be given, xargs will start the next task as soon as one of the currently running tasks finishes"; see `https://bluewaters.ncsa.illinois.edu/job-bundling` for more information.

```
#!/bin/bash
#PBS -A bakr
#PBS -j oe
#PBS -l nodes=1:ppn=16:xe
#PBS -l walltime=06:00:00
#PBS -l flags=commlocal:commtolerant

ts=$(date +%s)

source /opt/modules/default/init/bash

modl=$arg1
repl=$arg2

cd $PBS_O_WORKDIR

aprun -n 1 -N 1 -d 16 xargs -d '\n' -I cmd -P 16 /bin/bash -c 'cmd' <<EOF
./e_run_fasttree_for_timings.sh $modl $repl 1 62
./e_run_fasttree_for_timings.sh $modl $repl 63 125
./e_run_fasttree_for_timings.sh $modl $repl 126 187
./e_run_fasttree_for_timings.sh $modl $repl 188 250
./e_run_fasttree_for_timings.sh $modl $repl 251 312
./e_run_fasttree_for_timings.sh $modl $repl 313 375
./e_run_fasttree_for_timings.sh $modl $repl 376 437
./e_run_fasttree_for_timings.sh $modl $repl 438 500
./e_run_fasttree_for_timings.sh $modl $repl 401 562
./e_run_fasttree_for_timings.sh $modl $repl 563 625
./e_run_fasttree_for_timings.sh $modl $repl 626 687
./e_run_fasttree_for_timings.sh $modl $repl 688 750
./e_run_fasttree_for_timings.sh $modl $repl 751 812
./e_run_fasttree_for_timings.sh $modl $repl 813 875
./e_run_fasttree_for_timings.sh $modl $repl 876 937
./e_run_fasttree_for_timings.sh $modl $repl 938 1000
EOF

te=$(date +%s)
rt=$[te - ts]
echo "Estimated gene trees on $modl (exons) in $rt seconds"
```

Note that `./e_run_fasttree_for_timings.sh` simply runs FastTree-2 on genes numbered `$sgen` to `$egen`.

```
for gene in ‘seq -f "%04g" $sgen $egen‘; do
    $fasttree -nt \
              -gtr \
              -quiet \
              -log fasttree-$gene.log \
              $gene.phy > fasttree-$gene.tre
done
```

Note that FastTree-2 was compiled using the following commands:

```
module unload darshan
module swap PrgEnv-cray PrgEnv-gnu
module swp gcc gcc/5.2.0  # Target: x86_64-suse-linux
gcc -DUSE_DOUBLE -O3 -finline-functions -funroll-loops \
    -Wall -o FastTreeDbl-no-darshan FastTree.c -lm
```

Table S4: **Running times for gene tree estimation (100-taxon datasets).** Running times (in seconds) for gene tree estimation is the time required to run FastTree-2 on datasets with 100 taxa and either 1000 exons or 1000 introns using a single Blue Waters compute node with 64 GB of memory and 16 floating-point cores. On average ($\pm$ standard deviation), gene tree estimation required $1080 \pm 107$ seconds ($18 \pm 2$ minutes).

| Species Tree Height | Replicate Number | Time (s) *for 1000 exons* | Time (s) *for 1000 introns* |
|---|---|---|---|
| 10M | 01 | 1075 | 1074 |
| 10M | 02 | 884 | 946 |
| 10M | 03 | 1166 | 1264 |
| 10M | 04 | 1101 | 1095 |
| 10M | 05 | 1217 | 1097 |
| 10M | 06 | 1132 | 1121 |
| 10M | 07 | 1189 | 1361 |
| 10M | 08 | 1011 | 1042 |
| 10M | 09 | 1234 | 1241 |
| 10M | 10 | 1032 | 995 |
| 10M | 11 | 1113 | 1153 |
| 10M | 12 | 1084 | 1104 |
| 10M | 13 | 1127 | 1130 |
| 10M | 14 | 1091 | 1157 |
| 10M | 15 | 1218 | 1235 |
| 10M | 16 | 1106 | 1021 |
| 10M | 17 | 1173 | 1252 |
| 10M | 18 | 1111 | 1236 |
| 10M | 19 | 1167 | 1175 |
| 10M | 20 | 1060 | 1093 |
| 500K | 01 | 913 | 874 |
| 500K | 02 | 1172 | 1064 |
| 500K | 03 | 1126 | 1149 |
| 500K | 04 | 1025 | 1016 |
| 500K | 05 | 1014 | 1015 |
| 500K | 06 | 1000 | 1054 |
| 500K | 07 | 927 | 893 |
| 500K | 08 | 978 | 875 |
| 500K | 09 | 1181 | 1197 |
| 500K | 10 | 1093 | 1098 |
| 500K | 11 | 1042 | 995 |
| 500K | 12 | 987 | 904 |
| 500K | 13 | 1082 | 1015 |
| 500K | 14 | 964 | 941 |
| 500K | 15 | 844 | 835 |
| 500K | 16 | 1126 | 1126 |
| 500K | 17 | 1180 | 1104 |
| 500K | 18 | 1136 | 1113 |
| 500K | 19 | 1107 | 1066 |
| 500K | 20 | 1067 | 986 |

Table S5: **Running times for gene tree estimation (1000-taxon datasets).** Running time for gene tree estimation is the time required to run FastTree-2 on datasets with 1000 taxa and either 1000 exons or 1000 introns using a single Blue Waters compute node with 64 GB of memory and 16 floating-point cores. On average ($\pm$ standard deviation), gene tree estimation required $13018 \pm 1169$ seconds ($217 \pm 20$ minutes).

| Species Tree Height | Replicate Number | Time (s) *for 1000 exons* | Time (s) *for 1000 introns* |
|---|---|---|---|
| 10M | 01 | 13083 | 12295 |
| 10M | 02 | 13310 | 13014 |
| 10M | 03 | 10546 | 10948 |
| 10M | 04 | 12544 | 11781 |
| 10M | 05 | 13530 | 13673 |
| 10M | 06 | 12430 | 10969 |
| 10M | 07 | 14043 | 13735 |
| 10M | 08 | 14588 | 15273 |
| 10M | 09 | 13415 | 13045 |
| 10M | 10 | 14336 | 13740 |
| 10M | 11 | 14455 | 14214 |
| 10M | 12 | 12596 | 11266 |
| 10M | 13 | 13340 | 13056 |
| 10M | 14 | 12159 | 11721 |
| 10M | 15 | 14154 | 13084 |
| 10M | 16 | 13247 | 12422 |
| 10M | 17 | 13677 | 12949 |
| 10M | 18 | 11600 | 11295 |
| 10M | 19 | 13747 | 13045 |
| 10M | 20 | 13940 | 13176 |
| 500K | 01 | 11632 | 12232 |
| 500K | 02 | 11380 | 10640 |
| 500K | 03 | 13387 | 12627 |
| 500K | 04 | 13938 | 13733 |
| 500K | 05 | 12425 | 12280 |
| 500K | 06 | 9460 | 9648 |
| 500K | 07 | 13652 | 13085 |
| 500K | 08 | 12464 | 11671 |
| 500K | 09 | 13755 | 12320 |
| 500K | 10 | 13912 | 13785 |
| 500K | 11 | 14128 | 13534 |
| 500K | 12 | 14416 | 14221 |
| 500K | 13 | 12361 | 13029 |
| 500K | 14 | 14402 | 14394 |
| 500K | 15 | 13128 | 13920 |
| 500K | 16 | 14148 | 13238 |
| 500K | 17 | 14407 | 13344 |
| 500K | 18 | 14279 | 13794 |
| 500K | 19 | 12874 | 12264 |
| 500K | 20 | 14283 | 13778 |

#### Supplementary Results

We often specify the input parameters when referring to Neighbor Joining (NJ) and NJMerge. For example, NJ($D_{AGID}$) refers to NJ given the average gene tree internode distance (AGID) matrix as input, and NJMerge($\mathcal{T}_{RAX}, D_{AGID}$) refers to NJMerge given the constraint trees estimated using RAxML and the AGID matrix as input.

Distance matrices were created using two different approaches.

- $D_{AGID}$ refers to the Average Gene tree Internode Distance matrix (as described in [6]) from estimated gene trees using ASTRID [15].
- $D_{LD}$ refers to the logdet distance matrix (as described in [12]) and computed from concatenated alignment using PAUP* [14].

Constraint trees were created using four different approaches.

- $\mathcal{T}_{true}$ refers to constraint trees computed by restricting the true species tree to each subset of species.
- $\mathcal{T}_{AST}$ refers to constraint trees computed by running ASTRAL-III on each subset, i.e., on estimated gene trees restricted to subsets of species.
- $\mathcal{T}_{SVD}$ refers to constraint trees computed by running SVDquartets on each subset, i.e., on the concatenated alignment restricted to subsets of species.
- $\mathcal{T}_{RAX}$ refers to constraint trees computed by running RAxML on each subset, i.e., on the concatenated alignment restricted to subsets of species.

Table S6: **Method Failures.** Methods were run on 20 replicate datasets for each model condition with 1000 species, 1000 genes, two levels of ILS (species tree heights: 10M and 500K), and two sequence types (exon and intron). All four methods (ASTRAL-III, SVDquartets, RAxML, and NJMerge) failed on some datasets, as recorded below. ASTRAL-III failed due to running beyond the maximum wall-clock time of 48 hours; SVDquartets failed due to segmentation faults; RAxML failed due to running out of memory, and NJMerge failed due to being unable to find a legal siblinghood.

| # of Taxa | # of Genes | Species Tree Height | Data Type | Method | Fraction of Replicates | Replicate Numbers |
|---|---|---|---|---|---|---|
| 100 | 25 | 500K | exon | NJMerge($\mathcal{T}_{true}$, $D_{LD}$) | 1/20 | 10 |
| 100 | 25 | 500K | exon | NJMerge($\mathcal{T}_{RAX}$, $D_{AGID}$) | 1/20 | 16 |
| 100 | 25 | 500K | intron | NJMerge($\mathcal{T}_{true}$, $D_{AGID}$) | 1/20 | 6 |
| 1000 | 1000 | 10M | exon | SVDquartets | 20/20 | All |
| 1000 | 1000 | 10M | exon | RAxML | 3/20 | 2, 8, 17 |
| 1000 | 1000 | 10M | intron | NJMerge($\mathcal{T}_{AST}$, $D_{LD}$) | 1/20 | 20 |
| 1000 | 1000 | 10M | intron | SVDquartets | 20/20 | All |
| 1000 | 1000 | 10M | intron | RAxML | 20/20 | All |
| 1000 | 1000 | 500K | exon | ASTRAL-III | 19/20 | All except 15 |
| 1000 | 1000 | 500K | exon | NJMerge($\mathcal{T}_{true}$, $D_{LD}$) | 1/20 | 18 |
| 1000 | 1000 | 500K | exon | NJMerge($\mathcal{T}_{AST}$, $D_{LD}$) | 1/20 | 18 |
| 1000 | 1000 | 500K | exon | NJMerge($\mathcal{T}_{SVD}$, $D_{LD}$) | 2/20 | 14, 18 |
| 1000 | 1000 | 500K | exon | NJMerge($\mathcal{T}_{RAX}$, $D_{LD}$) | 2/20 | 14, 18 |
| 1000 | 1000 | 500K | exon | SVDquartets | 20/20 | All |
| 1000 | 1000 | 500K | intron | ASTRAL-III | 4/20 | 1, 5, 6, 20 |
| 1000 | 1000 | 500K | intron | NJMerge($\mathcal{T}_{SVD}, D_{LD}$) | 1/20 | 6 |
| 1000 | 1000 | 500K | intron | SVDquartets | 20/20 | All |
| 1000 | 1000 | 500K | intron | RAxML | 19/20 | All except 6 |

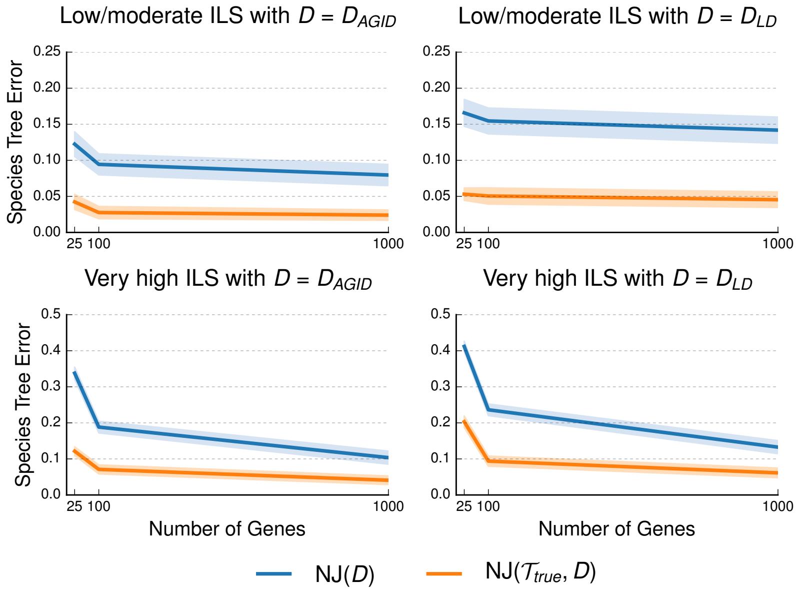

Figure S1: **Impact of distance matrix on NJMerge for 100-taxon, exon-like datasets.** Neighbor Joining (NJ) was run with two different distance matrices, and NJMerge was run with two different distance matrices and constraint trees that agreed with the true species tree (see the Performance Study section for more information on the notation). Datasets had two different levels of incomplete lineage sorting (ILS) and numbers of genes varying from 25 to 1000. Species tree estimation error is defined as the normalized Robinson-Foulds (RF) distance between true and estimated species trees. Lines represent the average over replicate datasets, and filled regions indicate the standard error.

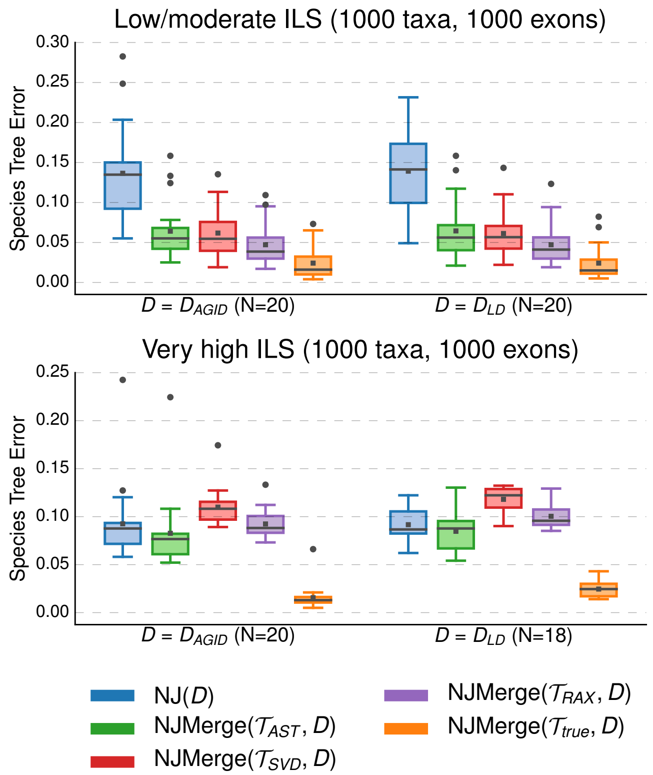

Figure S2: **Impact of constraint trees on NJMerge for 1000-taxon, exon-like datasets.** Neighbor Joining (NJ) was run with two different distance matrices, and NJMerge was run with two difference distance matrices and four different sets of constraint trees; see notation section above for details. Species tree estimation error is defined as the normalized Robinson-Foulds (RF) distance between true and estimated species trees. Note that gray bars represent medians, gray squares represent means, gray circles represent outliers, box plots are defined by quartiles (extending from the first to the third quartiles), and whiskers extend to plus/minus 1.5 times the interquartile distance (unless greater/less than the maximum/minimum value).

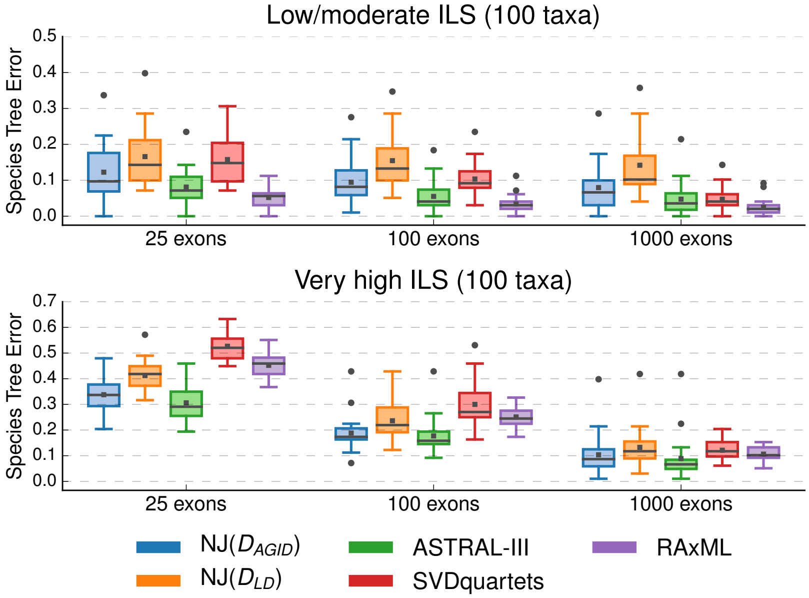

Figure S3: **Comparison of species tree methods for 100-taxon, exon-like datasets.** All methods were run on the full dataset (i.e., not subsets) with 100 species. Neighbor Joining (NJ) was run with two different distance matrices; see notation section above for details. Species tree estimation error is defined as the normalized Robinson-Foulds (RF) distance between true and estimated species trees. Note that gray bars represent medians, gray squares represent means, gray circles represent outliers, box plots are defined by quartiles (extending from the first to the third quartiles), and whiskers extend to plus/minus 1.5 times the interquartile distance (unless greater/less than the maximum/minimum value).

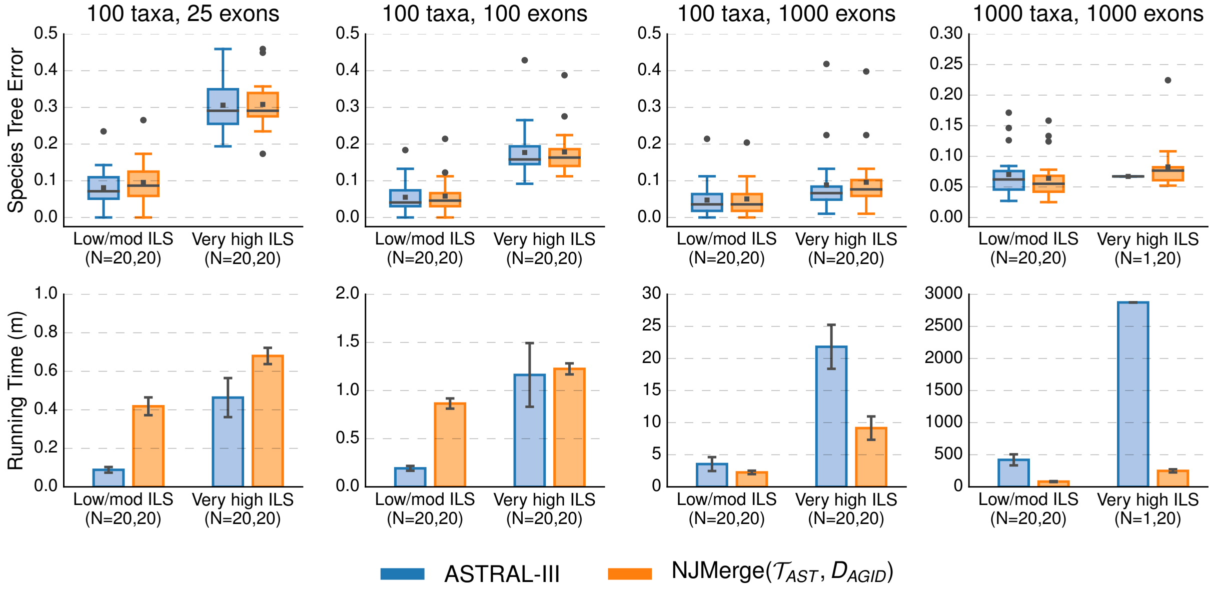

Figure S4: **Comparison of ASTRAL-III and NJMerge given ASTRAL-III constraint trees and AGID matrix for exon-like datasets.** Subplots on top row show species tree estimation error (defined as the normalized RF distance between true and estimated species trees); note that gray bars represent medians, gray squares represent means, gray circles represent outliers, box plots are defined by quartiles (extending from the first to the third quartiles), and whiskers extend to plus/minus 1.5 times the interquartile distance (unless greater/less than the maximum/minimum value). Subplots on bottom row show running time (in minutes); bars represent means and error bars represent standard deviations across replicate datasets. NJMerge running times is for computing the subset trees "in serial"; see Equation (1) in the main text for more information. The numbers of replicates on which the methods completed is shown on the x-axis, e.g., $N = X, Y$ indicates that ASTRAL-III completed on $X$ out of 20 replicates and that NJMerge($\mathcal{T}_{AST}, D_{LD}$) completed on $Y$ out of 20 replicates.

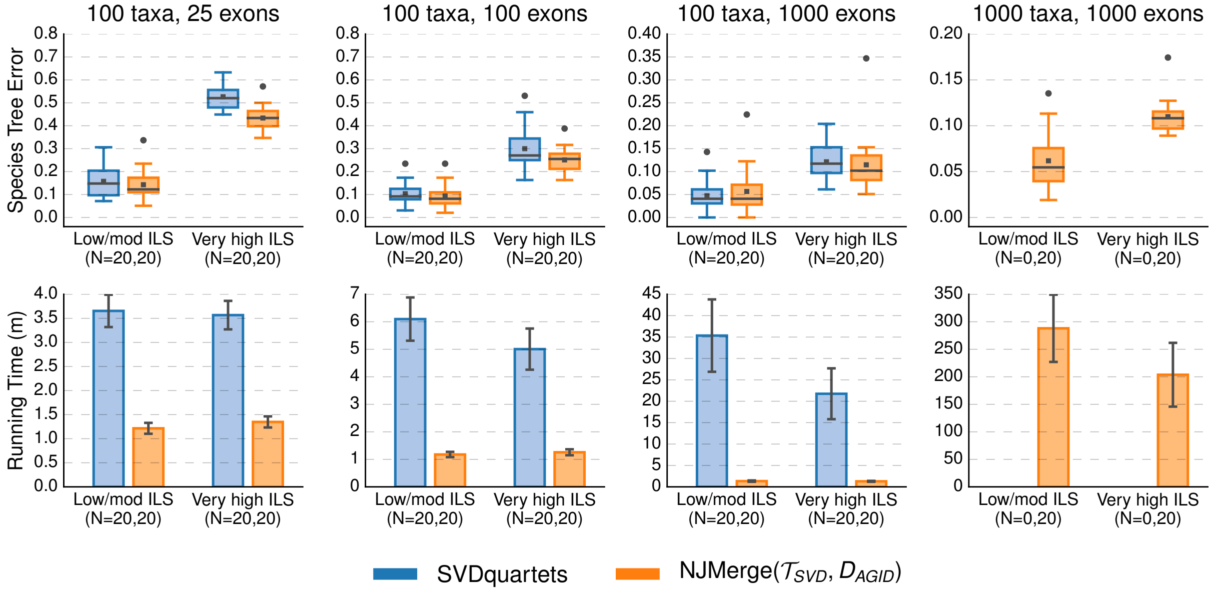

Figure S5: **Comparison of SVDquartets and NJMerge (given SVDquartets constraint trees and AGID matrix) for exon-like datasets.** Subplots on top row show species tree estimation error (defined as the normalized RF distance between true and estimated species trees); note that gray bars represent medians, gray squares represent means, gray circles represent outliers, box plots are defined by quartiles (extending from the first to the third quartiles), and whiskers extend to plus/minus 1.5 times the interquartile distance (unless greater/less than the maximum/minimum value). Subplots on bottom row show running time (in minutes); bars represent means and error bars represent standard deviations across replicate datasets. NJMerge running times is for computing the subset trees "in serial"; see Equation (1) in the main text for more information. The numbers of replicates on which the methods completed is shown on the x-axis, e.g., $N = X, Y$ indicates that SVDquartets completed on $X$ out of 20 replicates and that NJMerge($\mathcal{T}_{SVD}, D_{LD}$) completed on $Y$ out of 20 replicates.

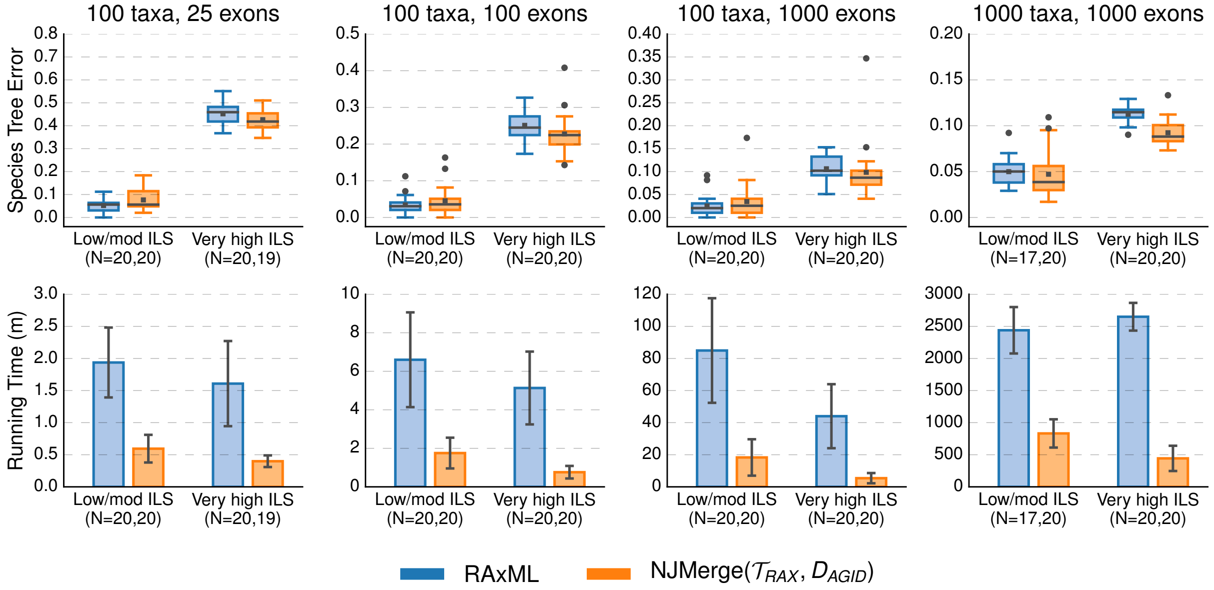

Figure S6: **Comparison of RAxML and NJMerge given RAxML constraint trees and AGID matrix for exon-like datasets.** Subplots on top row show species tree estimation error (defined as the normalized RF distance between true and estimated species trees); note that gray bars represent medians, gray squares represent means, gray circles represent outliers, box plots are defined by quartiles (extending from the first to the third quartiles), and whiskers extend to plus/minus 1.5 times the interquartile distance (unless greater/less than the maximum/minimum value). Subplots on bottom row show running time (in minutes); bars represent means and error bars represent standard deviations across replicate datasets. NJMerge running times is for computing the subset trees "in serial"; see Equation (1) in the main text for more information. The numbers of replicates on which the methods completed is shown on the x-axis, e.g., $N = X, Y$ indicates that RAxML completed on $X$ out of 20 replicates and that NJMerge($\mathcal{T}_{RAX}, D_{LD}$) completed on $Y$ out of 20 replicates.

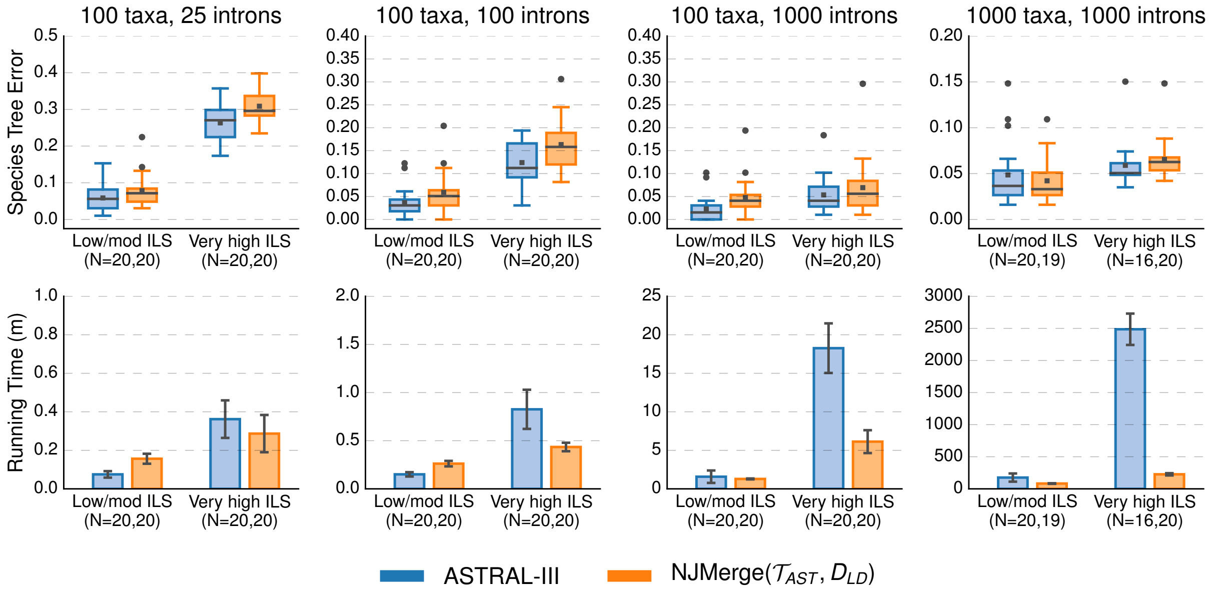

Figure S7: **Comparison of ASTAL-III and NJMerge given ASTRAL-III constraint trees and log-det distance matrix for intron-like datasets.** Subplots on top row show species tree estimation error (defined as the normalized RF distance between true and estimated species trees); note that gray bars represent medians, gray squares represent means, gray circles represent outliers, box plots are defined by quartiles (extending from the first to the third quartiles), and whiskers extend to plus/minus 1.5 times the interquartile distance (unless greater/less than the maximum/minimum value). Subplots on bottom row show running time (in minutes); bars represent means and error bars represent standard deviations across replicate datasets. NJMerge running times is for computing the subset trees "in serial"; see Equation (1) in the main text for more information. The numbers of replicates on which the methods completed is shown on the x-axis, e.g., $N = X, Y$ indicates that ASTRAL-III completed on $X$ out of 20 replicates and that NJMerge($\mathcal{T}_{AST}, D_{LD}$) completed on $Y$ out of 20 replicates.

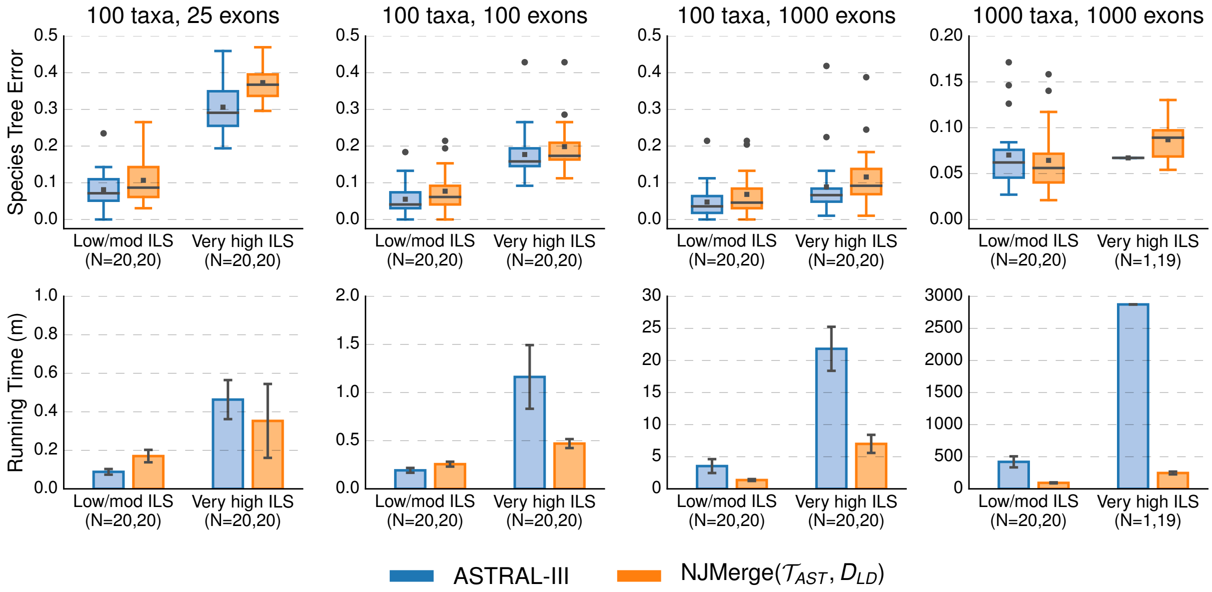

Figure S8: **Comparison of ASTRAL-III and NJMerge given ASTRAL-III constraint trees and log-det distance matrix for exon-like datasets.** Subplots on top row show species tree estimation error (defined as the normalized RF distance between true and estimated species trees); note that gray bars represent medians, gray squares represent means, gray circles represent outliers, box plots are defined by quartiles (extending from the first to the third quartiles), and whiskers extend to plus/minus 1.5 times the interquartile distance (unless greater/less than the maximum/minimum value). Subplots on bottom row show running time (in minutes); bars represent means and error bars represent standard deviations across replicate datasets. NJMerge running times is for computing the subset trees "in serial"; see Equation (1) in the main text for more information. The numbers of replicates on which the methods completed is shown on the x-axis, e.g., $N = X, Y$ indicates that ASTRAL-III completed on $X$ out of 20 replicates and that NJMerge($\mathcal{T}_{AST}, D_{LD}$) completed on $Y$ out of 20 replicates.

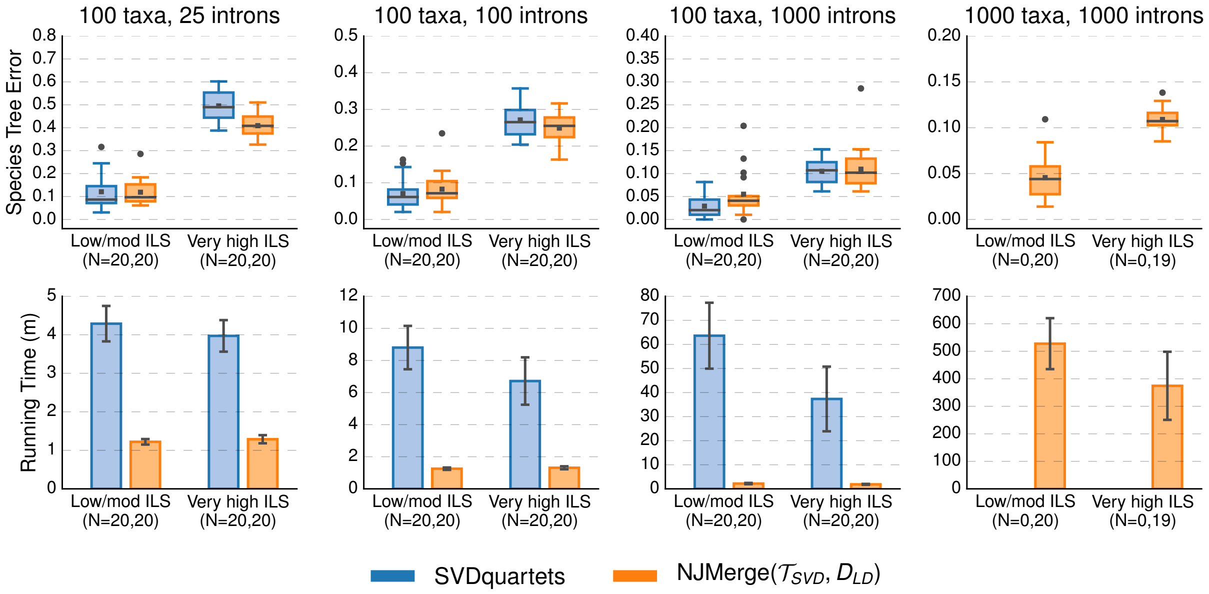

Figure S9: **Comparison of SVDquartets and NJMerge given SVDquartet constraint trees and log-det distance matrix for intron-like datasets.** Subplots on top row show species tree estimation error (defined as the normalized RF distance between true and estimated species trees); note that gray bars represent medians,, gray squares represent means, gray circles represent outliers, box plots are defined by quartiles (extending from the first to the third quartiles), and whiskers extend to plus/minus 1.5 times the interquartile distance (unless greater/less than the maximum/minimum value). Subplots on bottom row show running time (in minutes); bars represent means and error bars represent standard deviations across replicate datasets. NJMerge running times is for computing the subset trees "in serial"; see Equation (1) in the main text for more information. The numbers of replicates on which the methods completed is shown on the x-axis, e.g., $N = X, Y$ indicates that SVDquartets completed on $X$ out of 20 replicates and that NJMerge($\mathcal{T}_{SVD}, D_{LD}$) completed on $Y$ out of 20 replicates.

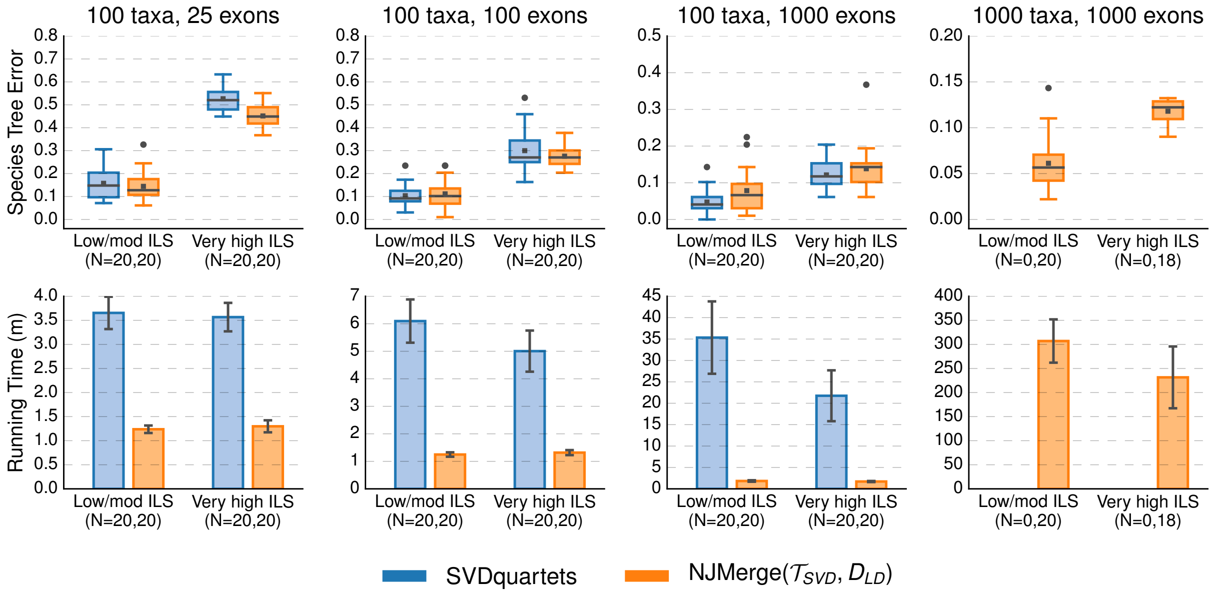

Figure S10: **Comparison of SVDquartets and NJMerge given log-det distance matrix for exon-like datasets.** Subplots on top row show species tree estimation error (defined as the normalized RF distance between true and estimated species trees); note that gray bars represent medians, gray squares represent means, gray circles represent outliers, box plots are defined by quartiles (extending from the first to the third quartiles), and whiskers extend to plus/minus 1.5 times the interquartile distance (unless greater/less than the maximum/minimum value). Subplots on bottom row show running time (in minutes); bars represent means and error bars represent standard deviations across replicate datasets. NJMerge running times is for computing the subset trees "in serial"; see Equation (1) in the main text for more information. The numbers of replicates on which the methods completed is shown on the x-axis, e.g., $N = X, Y$ indicates that SVDquartets completed on $X$ out of 20 replicates and that NJMerge($\mathcal{T}_{SVD}, D_{LD}$) completed on $Y$ out of 20 replicates.

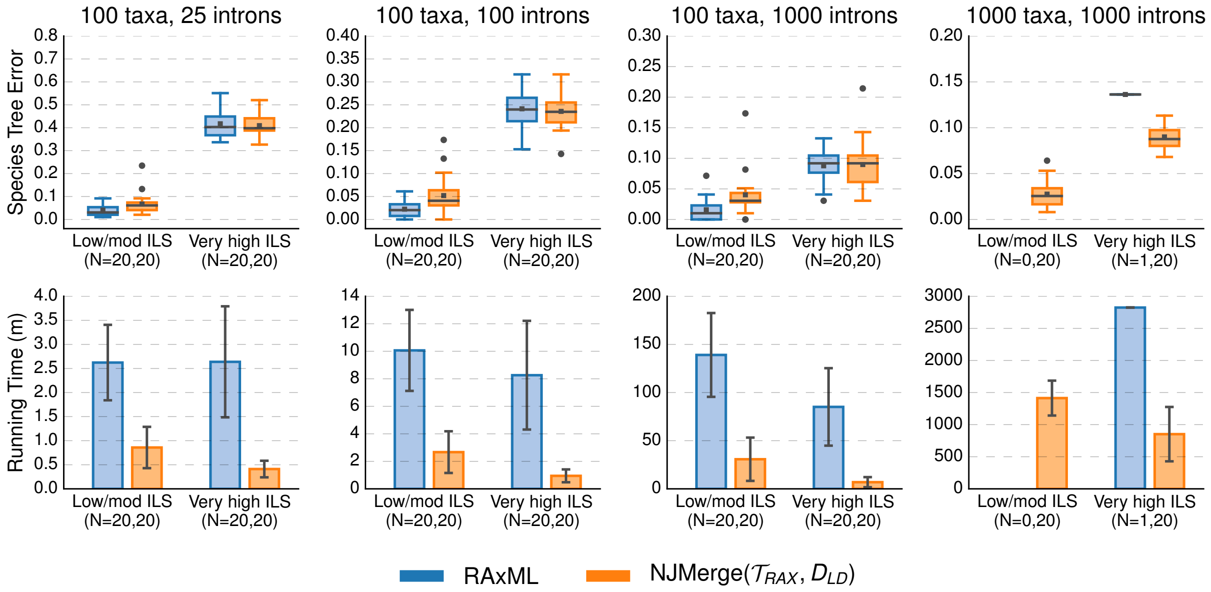

Figure S11: **Comparison of RAxML and NJMerge given RAxML constraint trees and log-det distance matrix for intron-like datasets.** Subplots on top row show species tree estimation error (defined as the normalized RF distance between true and estimated species trees); note that gray bars represent medians, gray squares represent means, gray circles represent outliers, box plots are defined by quartiles (extending from the first to the third quartiles), and whiskers extend to plus/minus 1.5 times the interquartile distance (unless greater/less than the maximum/minimum value). Subplots on bottom row show running time (in minutes); bars represent means and error bars represent standard deviations across replicate datasets. NJMerge running times is for computing the subset trees "in serial"; see Equation (1) in the main text for more information. The numbers of replicates on which the methods completed is shown on the x-axis, e.g., $N = X, Y$ indicates that RAxML completed on $X$ out of 20 replicates and that NJMerge($\mathcal{T}_{RAX}, D_{LD}$) completed on $Y$ out of 20 replicates.

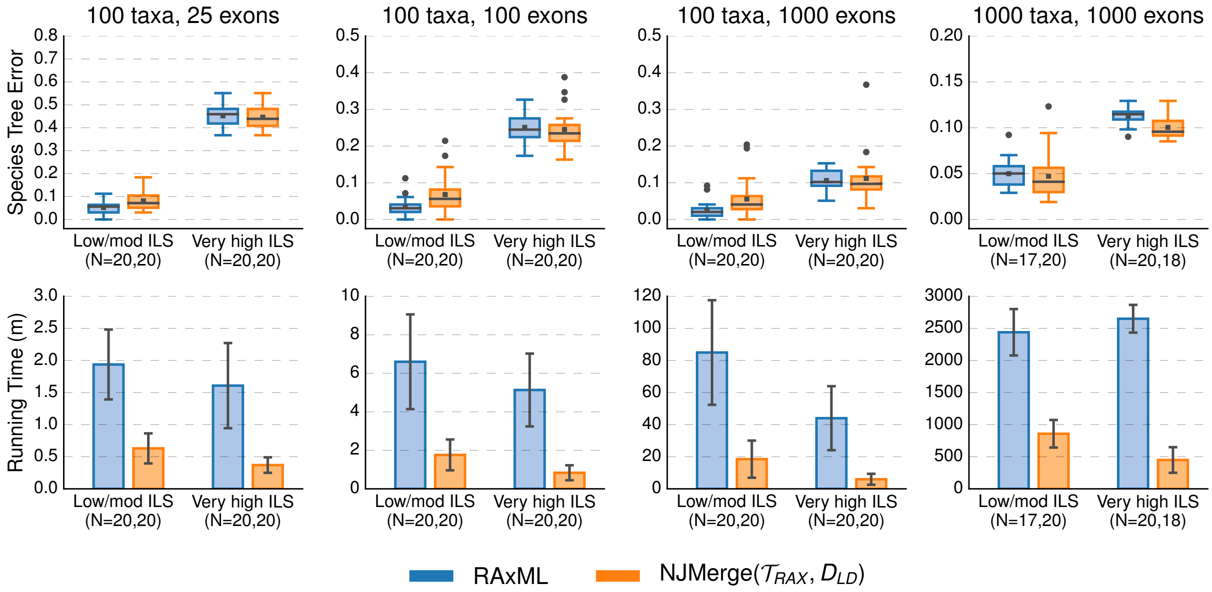

Figure S12: **Comparison of RAxML and NJMerge given RAxML constraint trees and log-det distance matrix for exon-like datasets.** Subplots on top row show species tree estimation error (defined as the normalized RF distance between true and estimated species trees); note that gray bars represent medians, gray squares represent means, gray circles represent outliers, box plots are defined by quartiles (extending from the first to the third quartiles), and whiskers extend to plus/minus 1.5 times the interquartile distance (unless greater/less than the maximum/minimum value). Subplots on bottom row show running time (in minutes); bars represent means and error bars represent standard deviations across replicate datasets. NJMerge running times is for computing the subset trees "in serial"; see Equation (1) in the main text for more information. The numbers of replicates on which the methods completed is shown on the x-axis, e.g., $N = X, Y$ indicates that RAxML completed on $X$ out of 20 replicates and that NJMerge($\mathcal{T}_{RAX}, D_{LD}$) completed on $Y$ out of 20 replicates.

Table S7: **Species tree error for NJMerge given AGID matrix.** Each species tree estimation method (ASTRAL-III, SVDquartets, or RAML) was run on the full dataset or on subsets in order to build constraint trees for NJMerge. We report the average (± standard deviation) species tree estimation error for *1)* the tree produced by running species tree method $M$ on the full set of species (all 100 or all 1000 taxa), *2)* the tree produced by running species tree method $M$ on subsets of species to produce $\mathcal{T}_M$, *3)* the tree produced by running NJ($D_{AGID}$), and *4)* running NJMerge($\mathcal{T}_M, D_{AGID}$). Species tree estimation error (defined as normalized RF distance between the true and the estimated species tree) was averaged across 20 replicate datasets, unless the number of replicate datasets is otherwise noted in parentheses. When methods were run on subsets, species tree estimation error was averaged across all subsets and all replicate datasets. Note that the number of taxa in the subset trees was less than 30 for the 100-taxon datasets and less than 120 for the 1000-taxon datasets.

| Number of Taxa | Number of Genes | Species Tree Height | Data Type | $M$ on full dataset | $M$ on subsets | NJ($D_{AGID}$) | NJMerge($\mathcal{T}_M, D_{AGID}$) |
|---|---|---|---|---|---|---|---|
| $M$ = *ASTRAL-III* | | | | | | | |
| 100 | 25 | 10M | exon | 0.08 | 0.07 | 0.12 | 0.10 |
| 100 | 25 | 10M | intron | 0.06 | 0.05 | 0.07 | 0.06 |
| 100 | 25 | 500K | exon | 0.31 | 0.25 | 0.34 | 0.31 |
| 100 | 25 | 500K | intron | 0.26 | 0.20 | 0.30 | 0.26 |
| 100 | 100 | 10M | exon | 0.06 | 0.04 | 0.09 | 0.06 |
| 100 | 100 | 10M | intron | 0.04 | 0.03 | 0.05 | 0.04 |
| 100 | 100 | 500K | exon | 0.18 | 0.12 | 0.19 | 0.18 |
| 100 | 100 | 500K | intron | 0.12 | 0.09 | 0.14 | 0.13 |
| 100 | 1000 | 10M | exon | 0.05 | 0.03 | 0.08 | 0.05 |
| 100 | 1000 | 10M | intron | 0.02 | 0.02 | 0.04 | 0.03 |
| 100 | 1000 | 500K | exon | 0.09 | 0.06 | 0.10 | 0.10 |
| 100 | 1000 | 500K | intron | 0.05 | 0.04 | 0.06 | 0.06 |
| 1000 | 1000 | 10M | exon | 0.07 | 0.04 | 0.14 | 0.06 |
| 1000 | 1000 | 10M | intron | 0.05 | 0.03 | 0.11 | 0.05 |
| 1000 | 1000 | 500K | exon | 0.07 (1) | 0.07 | 0.09 | 0.08 |
| 1000 | 1000 | 500K | intron | 0.06 (16) | 0.05 | 0.07 | 0.06 |
| $M$ = *SVDquartets* | | | | | | | |
| 100 | 25 | 10M | exon | 0.16 | 0.12 | 0.12 | 0.14 |
| 100 | 25 | 10M | intron | 0.12 | 0.09 | 0.07 | 0.10 |
| 100 | 25 | 500K | exon | 0.53 | 0.38 | 0.34 | 0.43 |
| 100 | 25 | 500K | intron | 0.50 | 0.34 | 0.30 | 0.39 |
| 100 | 100 | 10M | exon | 0.10 | 0.08 | 0.09 | 0.09 |
| 100 | 100 | 10M | intron | 0.07 | 0.06 | 0.05 | 0.07 |
| 100 | 100 | 500K | exon | 0.30 | 0.21 | 0.19 | 0.25 |
| 100 | 100 | 500K | intron | 0.27 | 0.20 | 0.14 | 0.23 |
| 100 | 1000 | 10M | exon | 0.05 | 0.04 | 0.08 | 0.06 |
| 100 | 1000 | 10M | intron | 0.03 | 0.03 | 0.04 | 0.03 |
| 100 | 1000 | 500K | exon | 0.12 | 0.08 | 0.10 | 0.11 |
| 100 | 1000 | 500K | intron | 0.11 | 0.08 | 0.06 | 0.10 |
| 1000 | 1000 | 10M | exon | NA (0) | 0.04 | 0.14 | 0.06 |
| 1000 | 1000 | 10M | intron | NA (0) | 0.03 | 0.11 | 0.05 |
| 1000 | 1000 | 500K | exon | NA (0) | 0.10 | 0.09 | 0.11 |
| 1000 | 1000 | 500K | intron | NA (0) | 0.09 | 0.07 | 0.10 |
| $M$ = *RAxML* | | | | | | | |
| 100 | 25 | 10M | exon | 0.05 | 0.04 | 0.12 | 0.08 |
| 100 | 25 | 10M | intron | 0.04 | 0.03 | 0.07 | 0.05 |
| 100 | 25 | 500K | exon | 0.45 | 0.38 | 0.34 | 0.43 (19) |
| 100 | 25 | 500K | intron | 0.42 | 0.34 | 0.30 | 0.39 |
| 100 | 100 | 10M | exon | 0.03 | 0.02 | 0.09 | 0.05 |
| 100 | 100 | 10M | intron | 0.02 | 0.02 | 0.05 | 0.03 |
| 100 | 100 | 500K | exon | 0.25 | 0.18 | 0.19 | 0.23 |
| 100 | 100 | 500K | intron | 0.24 | 0.18 | 0.14 | 0.21 |
| 100 | 1000 | 10M | exon | 0.02 | 0.01 | 0.08 | 0.03 |
| 100 | 1000 | 10M | intron | 0.02 | 0.01 | 0.04 | 0.02 |
| 100 | 1000 | 500K | exon | 0.11 | 0.06 | 0.10 | 0.10 |
| 100 | 1000 | 500K | intron | 0.09 | 0.06 | 0.06 | 0.08 |
| 1000 | 1000 | 10M | exon | 0.05 (17) | 0.02 | 0.14 | 0.05 |
| 1000 | 1000 | 10M | intron | NA (0) | 0.01 | 0.11 | 0.03 |
| 1000 | 1000 | 500K | exon | 0.11 | 0.08 | 0.09 | 0.09 |
| 1000 | 1000 | 500K | intron | 0.14 (1) | 0.08 | 0.07 | 0.08 |

Table S8: **Species tree error for NJMerge given log-det distance matrix.** Each species tree estimation method (ASTRAL-III, SVDquartets, or RAML) was run on the full dataset or on subsets in order to build constraint trees for NJMerge. We report the average (± standard deviation) species tree estimation error for *1)* the tree produced by running species tree method $M$ on the full set of species (all 100 or all 1000 taxa), *2)* the tree produced by running species tree method $M$ on subsets of species to produce $\mathcal{T}_M$, *3)* the tree produced by running NJ($D_{LD}$), and *4)* running NJMerge($\mathcal{T}_M, D_{LD}$). Species tree estimation error (defined as normalized RF distance between the true and the estimated species tree) was averaged across 20 replicate datasets, unless the number of replicate datasets is otherwise noted in parentheses. When methods were run on subsets, species tree estimation error was averaged across all subsets and all replicate datasets. Note that the number of taxa in the subset trees was less than 30 for the 100-taxon datasets and less than 120 for the 1000-taxon datasets.

| Number of Taxa | Number of Genes | Species Tree Height | Data Type | $M$ on full dataset | $M$ on subsets | NJ($D_{LD}$) | NJMerge($\mathcal{T}_M, D_{LD}$) |
|---|---|---|---|---|---|---|---|
| *$M$ = ASTRAL-III* | | | | | | | |
| 100 | 25 | 10M | exon | 0.08 | 0.06 | 0.17 | 0.11 |
| 100 | 25 | 10M | intron | 0.06 | 0.04 | 0.14 | 0.08 |
| 100 | 25 | 500K | exon | 0.31 | 0.26 | 0.41 | 0.37 |
| 100 | 25 | 500K | intron | 0.26 | 0.19 | 0.36 | 0.31 |
| 100 | 100 | 10M | exon | 0.06 | 0.04 | 0.15 | 0.08 |
| 100 | 100 | 10M | intron | 0.04 | 0.03 | 0.12 | 0.06 |
| 100 | 100 | 500K | exon | 0.18 | 0.13 | 0.24 | 0.20 |
| 100 | 100 | 500K | intron | 0.12 | 0.10 | 0.21 | 0.16 |
| 100 | 1000 | 10M | exon | 0.05 | 0.03 | 0.14 | 0.07 |
| 100 | 1000 | 10M | intron | 0.02 | 0.02 | 0.11 | 0.05 |
| 100 | 1000 | 500K | exon | 0.09 | 0.06 | 0.13 | 0.12 |
| 100 | 1000 | 500K | intron | 0.05 | 0.04 | 0.09 | 0.07 |
| 1000 | 1000 | 10M | exon | 0.07 | 0.04 | 0.14 | 0.06 |
| 1000 | 1000 | 10M | intron | 0.05 | 0.03 | 0.11 | 0.04 (19) |
| 1000 | 1000 | 500K | exon | 0.07 (1) | 0.07 | 0.10 | 0.09 (19) |
| 1000 | 1000 | 500K | intron | 0.06 (16) | 0.05 | 0.07 | 0.07 |
| *$M$ = SVDquartets* | | | | | | | |
| 100 | 25 | 10M | exon | 0.16 | 0.11 | 0.17 | 0.14 |
| 100 | 25 | 10M | intron | 0.12 | 0.09 | 0.14 | 0.12 |
| 100 | 25 | 500K | exon | 0.53 | 0.36 | 0.41 | 0.45 |
| 100 | 25 | 500K | intron | 0.50 | 0.32 | 0.36 | 0.41 |
| 100 | 100 | 10M | exon | 0.10 | 0.07 | 0.15 | 0.11 |
| 100 | 100 | 10M | intron | 0.07 | 0.05 | 0.12 | 0.08 |
| 100 | 100 | 500K | exon | 0.30 | 0.22 | 0.24 | 0.28 |
| 100 | 100 | 500K | intron | 0.27 | 0.19 | 0.21 | 0.25 |
| 100 | 1000 | 10M | exon | 0.05 | 0.04 | 0.14 | 0.08 |
| 100 | 1000 | 10M | intron | 0.03 | 0.03 | 0.11 | 0.06 |
| 100 | 1000 | 500K | exon | 0.12 | 0.09 | 0.13 | 0.14 |
| 100 | 1000 | 500K | intron | 0.11 | 0.08 | 0.09 | 0.11 |
| 1000 | 1000 | 10M | exon | NA (0) | 0.04 | 0.14 | 0.06 |
| 1000 | 1000 | 10M | intron | NA (0) | 0.03 | 0.11 | 0.05 |
| 1000 | 1000 | 500K | exon | NA (0) | 0.10 | 0.10 | 0.12 (18) |
| 1000 | 1000 | 500K | intron | NA (0) | 0.09 | 0.07 | 0.11 (19) |
| *$M$ = RAxML* | | | | | | | |
| 100 | 25 | 10M | exon | 0.05 | 0.04 | 0.17 | 0.08 |
| 100 | 25 | 10M | intron | 0.04 | 0.03 | 0.14 | 0.07 |
| 100 | 25 | 500K | exon | 0.45 | 0.34 | 0.41 | 0.45 |
| 100 | 25 | 500K | intron | 0.42 | 0.32 | 0.36 | 0.41 |
| 100 | 100 | 10M | exon | 0.03 | 0.02 | 0.15 | 0.07 |
| 100 | 100 | 10M | intron | 0.02 | 0.02 | 0.12 | 0.05 |
| 100 | 100 | 500K | exon | 0.25 | 0.19 | 0.24 | 0.25 |
| 100 | 100 | 500K | intron | 0.24 | 0.18 | 0.21 | 0.24 |
| 100 | 1000 | 10M | exon | 0.02 | 0.01 | 0.14 | 0.06 |
| 100 | 1000 | 10M | intron | 0.02 | 0.01 | 0.11 | 0.04 |
| 100 | 1000 | 500K | exon | 0.11 | 0.06 | 0.13 | 0.11 |
| 100 | 1000 | 500K | intron | 0.09 | 0.06 | 0.09 | 0.09 |
| 1000 | 1000 | 10M | exon | 0.05 (17) | 0.02 | 0.14 | 0.05 |
| 1000 | 1000 | 10M | intron | NA (0) | 0.01 | 0.11 | 0.03 |
| 1000 | 1000 | 500K | exon | 0.11 | 0.08 | 0.10 | 0.10 (18) |
| 1000 | 1000 | 500K | intron | 0.14 (1) | 0.07 | 0.07 | 0.09 |

Table S9: **Running times for NJMerge given AGID matrix.** Each species tree estimation method $M$ was run on the full dataset (all 100 or all 1000 taxa) or on subsets in order to build a set $\mathcal{T}_M$ of constraint trees for NJMerge. We report the average running time ($\pm$ the standard deviation) in **seconds** across 20 replicate datasets, unless the number of replicate datasets is otherwise noted in parentheses. When methods were run on subsets, the time was measured per subset, and then average was taken across all subsets for all replicate datasets. Note that the 100-taxon datasets were decomposed into 4-6 subsets with a maximum subset size of 30 taxa and that the 1000-taxon datasets were decomposed into 10-15 subsets with a maximum subset size of 120 taxa.

| Number of Taxa | Number of Genes | Species Tree Height | Data Type | $M$ on full dataset | $M$ on subsets | NJMerge($\mathcal{T}_M, D_{AGID}$) |
|---|---|---|---|---|---|---|
| $M$ = *ASTRAL-III* | | | | | | |
| 100 | 25 | 10M | exon | $5 \pm 1$ | $4 \pm 1$ | $5 \pm 2$ |
| 100 | 25 | 10M | intron | $5 \pm 1$ | $4 \pm 1$ | $5 \pm 1$ |
| 100 | 25 | 500K | exon | $28 \pm 6$ | $8 \pm 2$ | $5 \pm 1$ |
| 100 | 25 | 500K | intron | $22 \pm 6$ | $7 \pm 2$ | $6 \pm 2$ |
| 100 | 100 | 10M | exon | $12 \pm 2$ | $10 \pm 2$ | $5 \pm 1$ |
| 100 | 100 | 10M | intron | $9 \pm 1$ | $9 \pm 2$ | $5 \pm 1$ |
| 100 | 100 | 500K | exon | $70 \pm 20$ | $15 \pm 3$ | $5 \pm 1$ |
| 100 | 100 | 500K | intron | $50 \pm 12$ | $14 \pm 3$ | $5 \pm 1$ |
| 100 | 1000 | 10M | exon | $213 \pm 65$ | $28 \pm 10$ | $5 \pm 1$ |
| 100 | 1000 | 10M | intron | $95 \pm 48$ | $21 \pm 5$ | $5 \pm 1$ |
| 100 | 1000 | 500K | exon | $1309 \pm 206$ | $121 \pm 56$ | $5 \pm 1$ |
| 100 | 1000 | 500K | intron | $1096 \pm 193$ | $103 \pm 57$ | $5 \pm 1$ |
| 1000 | 1000 | 10M | exon | $25231 \pm 5154$ | $239 \pm 119$ | $1939 \pm 66$ |
| 1000 | 1000 | 10M | intron | $10545 \pm 3823$ | $126 \pm 68$ | $1939 \pm 74$ |
| 1000 | 1000 | 500K | exon | $172346 \pm 0$ (1) | $1073 \pm 529$ | $1950 \pm 283$ |
| 1000 | 1000 | 500K | intron | $149146 \pm 14657$ (16) | $907 \pm 394$ | $1879 \pm 24$ |
| $M$ = *SVDquartets* | | | | | | |
| 100 | 25 | 10M | exon | $219 \pm 20$ | $15 \pm 5$ | $6 \pm 2$ |
| 100 | 25 | 10M | intron | $257 \pm 28$ | $15 \pm 5$ | $5 \pm 2$ |
| 100 | 25 | 500K | exon | $214 \pm 18$ | $17 \pm 5$ | $9 \pm 6$ |
| 100 | 25 | 500K | intron | $238 \pm 25$ | $15 \pm 5$ | $8 \pm 4$ |
| 100 | 100 | 10M | exon | $366 \pm 47$ | $14 \pm 5$ | $5 \pm 1$ |
| 100 | 100 | 10M | intron | $528 \pm 81$ | $14 \pm 5$ | $6 \pm 2$ |
| 100 | 100 | 500K | exon | $300 \pm 45$ | $15 \pm 6$ | $6 \pm 2$ |
| 100 | 100 | 500K | intron | $403 \pm 89$ | $15 \pm 6$ | $6 \pm 2$ |
| 100 | 1000 | 10M | exon | $2120 \pm 507$ | $16 \pm 7$ | $5 \pm 2$ |
| 100 | 1000 | 10M | intron | $3817 \pm 821$ | $19 \pm 9$ | $5 \pm 2$ |
| 100 | 1000 | 500K | exon | $1305 \pm 356$ | $16 \pm 6$ | $5 \pm 2$ |
| 100 | 1000 | 500K | intron | $2240 \pm 806$ | $16 \pm 6$ | $5 \pm 2$ |
| 1000 | 1000 | 10M | exon | $NA \pm NA$ (0) | $1238 \pm 1142$ | $2005 \pm 124$ |
| 1000 | 1000 | 10M | intron | $NA \pm NA$ (0) | $2219 \pm 2019$ | $1999 \pm 184$ |
| 1000 | 1000 | 500K | exon | $NA \pm NA$ (0) | $839 \pm 803$ | $2057 \pm 178$ |
| 1000 | 1000 | 500K | intron | $NA \pm NA$ (0) | $1550 \pm 1615$ | $1975 \pm 76$ |
| $M$ = *RAxML* | | | | | | |
| 100 | 25 | 10M | exon | $116 \pm 33$ | $7 \pm 5$ | $5 \pm 2$ |
| 100 | 25 | 10M | intron | $157 \pm 47$ | $10 \pm 7$ | $5 \pm 2$ |
| 100 | 25 | 500K | exon | $96 \pm 40$ | $4 \pm 3$ | $8 \pm 2$ (19) |
| 100 | 25 | 500K | intron | $158 \pm 69$ | $4 \pm 3$ | $8 \pm 3$ |
| 100 | 100 | 10M | exon | $396 \pm 148$ | $22 \pm 16$ | $5 \pm 2$ |
| 100 | 100 | 10M | intron | $604 \pm 177$ | $32 \pm 25$ | $5 \pm 1$ |
| 100 | 100 | 500K | exon | $308 \pm 113$ | $9 \pm 7$ | $6 \pm 2$ |
| 100 | 100 | 500K | intron | $496 \pm 237$ | $10 \pm 9$ | $5 \pm 2$ |
| 100 | 1000 | 10M | exon | $5097 \pm 1955$ | $238 \pm 202$ | $5 \pm 1$ |
| 100 | 1000 | 10M | intron | $8343 \pm 2611$ | $426 \pm 401$ | $5 \pm 1$ |
| 100 | 1000 | 500K | exon | $2641 \pm 1196$ | $71 \pm 60$ | $5 \pm 1$ |
| 100 | 1000 | 500K | intron | $5106 \pm 2414$ | $83 \pm 99$ | $5 \pm 1$ |
| 1000 | 1000 | 10M | exon | $146329 \pm 21692$ (17) | $3887 \pm 2023$ | $2055 \pm 165$ |
| 1000 | 1000 | 10M | intron | $NA \pm NA$ (0) | $6496 \pm 3226$ | $2010 \pm 174$ |
| 1000 | 1000 | 500K | exon | $158973 \pm 12955$ | $2037 \pm 1554$ | $2006 \pm 179$ |
| 1000 | 1000 | 500K | intron | $169440 \pm 0$ (1) | $3976 \pm 3203$ | $1933 \pm 86$ |

Table S10: **Running times for NJMerge given log-det distance matrix.** Each species tree estimation method $M$ was run on the full dataset (all 100 or all 1000 taxa) or on subsets in order to build a set $\mathcal{T}_M$ of constraint trees for NJMerge. We report the average running time ($\pm$ the standard deviation) in **seconds** across 20 replicate datasets, unless the number of replicate datasets is otherwise noted in parentheses. When methods were run on subsets, the time was measured per subset, and then average was taken across all subsets for all replicate datasets. Note that the 100-taxon datasets were decomposed into 4-6 subsets with a maximum subset size of 30 taxa and that the 1000-taxon datasets were decomposed into 10-14 subsets with a maximum subset size of 120 taxa.

| Number of Taxa | Number of Genes | Species Tree Height | Data Type | $M$ on full dataset | $M$ on subsets | NJMerge($\mathcal{T}_M, D_{LD}$) |
|---|---|---|---|---|---|---|
| $M$ = *ASTRAL-III* | | | | | | |
| 100 | 25 | 10M | exon | $5 \pm 1$ | $1 \pm 0$ | $6 \pm 2$ |
| 100 | 25 | 10M | intron | $5 \pm 1$ | $1 \pm 0$ | $5 \pm 1$ |
| 100 | 25 | 500K | exon | $28 \pm 6$ | $2 \pm 1$ | $13 \pm 11$ |
| 100 | 25 | 500K | intron | $22 \pm 6$ | $2 \pm 0$ | $9 \pm 6$ |
| 100 | 100 | 10M | exon | $12 \pm 2$ | $2 \pm 0$ | $5 \pm 1$ |
| 100 | 100 | 10M | intron | $9 \pm 1$ | $1 \pm 0$ | $5 \pm 1$ |
| 100 | 100 | 500K | exon | $70 \pm 20$ | $4 \pm 2$ | $6 \pm 2$ |
| 100 | 100 | 500K | intron | $50 \pm 12$ | $4 \pm 1$ | $6 \pm 1$ |
| 100 | 1000 | 10M | exon | $213 \pm 65$ | $10 \pm 5$ | $5 \pm 2$ |
| 100 | 1000 | 10M | intron | $95 \pm 48$ | $6 \pm 2$ | $5 \pm 1$ |
| 100 | 1000 | 500K | exon | $1309 \pm 206$ | $86 \pm 44$ | $5 \pm 1$ |
| 100 | 1000 | 500K | intron | $1096 \pm 193$ | $73 \pm 40$ | $5 \pm 1$ |
| 1000 | 1000 | 10M | exon | $25231 \pm 5154$ | $166 \pm 80$ | $2117 \pm 309$ |
| 1000 | 1000 | 10M | intron | $10545 \pm 3823$ | $75 \pm 45$ | $2004 \pm 123$ (19) |
| 1000 | 1000 | 500K | exon | $172346 \pm 0$ (1) | $945 \pm 482$ | $2126 \pm 343$ (19) |
| 1000 | 1000 | 500K | intron | $149146 \pm 14657$ (16) | $773 \pm 364$ | $1991 \pm 128$ |
| $M$ = *SVDquartets* | | | | | | |
| 100 | 25 | 10M | exon | $219 \pm 20$ | $16 \pm 5$ | $5 \pm 2$ |
| 100 | 25 | 10M | intron | $257 \pm 28$ | $15 \pm 5$ | $5 \pm 2$ |
| 100 | 25 | 500K | exon | $214 \pm 18$ | $15 \pm 6$ | $7 \pm 3$ |
| 100 | 25 | 500K | intron | $238 \pm 25$ | $15 \pm 5$ | $7 \pm 3$ |
| 100 | 100 | 10M | exon | $366 \pm 47$ | $15 \pm 5$ | $5 \pm 1$ |
| 100 | 100 | 10M | intron | $528 \pm 81$ | $14 \pm 5$ | $5 \pm 1$ |
| 100 | 100 | 500K | exon | $300 \pm 45$ | $16 \pm 5$ | $6 \pm 2$ |
| 100 | 100 | 500K | intron | $403 \pm 89$ | $15 \pm 5$ | $6 \pm 2$ |
| 100 | 1000 | 10M | exon | $2120 \pm 507$ | $16 \pm 7$ | $5 \pm 1$ |
| 100 | 1000 | 10M | intron | $3817 \pm 821$ | $18 \pm 9$ | $5 \pm 1$ |
| 100 | 1000 | 500K | exon | $1305 \pm 356$ | $16 \pm 6$ | $5 \pm 1$ |
| 100 | 1000 | 500K | intron | $2240 \pm 806$ | $16 \pm 6$ | $5 \pm 1$ |
| 1000 | 1000 | 10M | exon | $NA \pm NA$ (0) | $1225 \pm 1044$ | $2064 \pm 324$ |
| 1000 | 1000 | 10M | intron | $NA \pm NA$ (0) | $2288 \pm 2003$ | $2022 \pm 202$ |
| 1000 | 1000 | 500K | exon | $NA \pm NA$ (0) | $858 \pm 845$ | $2283 \pm 457$ (18) |
| 1000 | 1000 | 500K | intron | $NA \pm NA$ (0) | $1459 \pm 1514$ | $2081 \pm 216$ (19) |
| $M$ = *RAxML* | | | | | | |
| 100 | 25 | 10M | exon | $116 \pm 33$ | $7 \pm 5$ | $5 \pm 1$ |
| 100 | 25 | 10M | intron | $157 \pm 47$ | $10 \pm 8$ | $5 \pm 1$ |
| 100 | 25 | 500K | exon | $96 \pm 40$ | $3 \pm 2$ | $7 \pm 4$ |
| 100 | 25 | 500K | intron | $158 \pm 69$ | $4 \pm 4$ | $6 \pm 1$ |
| 100 | 100 | 10M | exon | $396 \pm 148$ | $22 \pm 15$ | $5 \pm 1$ |
| 100 | 100 | 10M | intron | $604 \pm 177$ | $32 \pm 25$ | $5 \pm 1$ |
| 100 | 100 | 500K | exon | $308 \pm 113$ | $9 \pm 7$ | $6 \pm 1$ |
| 100 | 100 | 500K | intron | $496 \pm 237$ | $11 \pm 10$ | $6 \pm 2$ |
| 100 | 1000 | 10M | exon | $5097 \pm 1955$ | $231 \pm 201$ | $5 \pm 2$ |
| 100 | 1000 | 10M | intron | $8343 \pm 2611$ | $390 \pm 366$ | $5 \pm 2$ |
| 100 | 1000 | 500K | exon | $2641 \pm 1196$ | $73 \pm 63$ | $5 \pm 2$ |
| 100 | 1000 | 500K | intron | $5106 \pm 2414$ | $84 \pm 96$ | $5 \pm 2$ |
| 1000 | 1000 | 10M | exon | $146329 \pm 21692$ (17) | $3935 \pm 1945$ | $2200 \pm 485$ |
| 1000 | 1000 | 10M | intron | $NA \pm NA$ (0) | $6697 \pm 3093$ | $2070 \pm 232$ |
| 1000 | 1000 | 500K | exon | $158973 \pm 12955$ | $2022 \pm 1463$ | $2076 \pm 200$ (18) |
| 1000 | 1000 | 500K | intron | $169440 \pm 0$ (1) | $3905 \pm 3174$ | $2259 \pm 1164$ |
